# CCB79 is a primate-specific cilium initiation factor essential to maintain neural progenitor diversity in developing brain tissue

**DOI:** 10.1101/2025.08.01.668123

**Authors:** Dhanasekaran Rathinam, Vaibhav Jadhav, Seema Wasim, Dina Elkahwagy, Nazlican Altinisik, Enes Cicek, Tejas Borkar, Aruljothi Mariappan, Omkar Suhas Vinchure, Nadine Poempner, Ziliang Zhao, Samhan Alsolami, Mo Li, Johannes Ptok, Christian Eggeling, Muhammad Tariq, Maria Giovanna Riparbelli, Emilio Cirri, Giuliano Callaini, Peter Nürnberg, Elke Gabriel, Jay Gopalakrishnan

## Abstract

Identifying the genes that regulate the accurate spatiotemporal diversity of neural progenitor cells (NPCs) helps to understand the mechanisms of human neocortex expansion. In primate brains, an additional intermediate progenitor layer, the outer subventricular zone (oSVZ), facilitates the expansion of the neocortex. Here, we identify an uncharacterized gene, *KIAA0408,* and show that its expression is enriched in intermediate progenitors. Removing *KIAA0408* in human-induced pluripotent stem cell (iPSC)-derived brain organoids results in an impaired cortical organization characterized by abnormal cell fate and patterning defects, including the depletion of intermediate and ventral progenitors. Molecularly, *KIAA0408* codes for a 79-kilodalton centriolar distal appendage protein (DAP) that controls cilium biogenesis (hereafter CCB79). CCB79 forms a complex with other DAP components and specifically localizes in the DAP at the onset of ciliogenesis, and its absence blocks ciliogenesis. Mechanistically, progenitors in 3D brain tissues are unable to form cilia, which induces aberrant hedgehog signaling and causes premature differentiation. Finally, human CCB79, rather than the mouse ortholog, rescues cilia defects, suggesting that CCB79 has undergone rapid evolution from rodents to primates to fine-tune ciliogenesis for proper brain development.

## Introduction

Primate evolution is characterized by a significant expansion of the forebrain, resulting in a disproportionately large neocortex^1–5^. Emerging evidence indicates that these evolutionary differences originate at the NPC level—the fundamental unit of the developing brain^2,4–8^. NPC’s spatiotemporal diversity is carefully coordinated to promote the proper development of the cerebral cortex. In the developing brain, NPCs are located in distinct layers of the ventricular zone (VZ). Here, apical radial glia divide symmetrically to produce dorsal and ventral intermediate progenitors, which migrate to the subventricular zone (SVZ) and can proliferate further before symmetrically differentiating into migrating neurons across all cortical layers^9–12^. In species with more prominent, gyrencephalic brains (e.g., primates), an additional outer subventricular zone (oSVZ) contains a large number of outer radial glial cells and intermediate progenitors, a highly neurogenic NPC type that greatly promotes neocortex expansion^13–16^.

Conversely, rodents, which typically have smooth-surfaced (lissencephalic) brains, have fewer outer radial glial cells, intermediate progenitors, and a less distinct oSVZ^13,16^. The surface area and thickness of the cerebral cortex in primates are much greater than in rodents due to the increased diversity of NPCs and neurons^1–5^. This wide range of NPC types and their developmental niches shows how the balance between self-renewing divisions and differentiation can fundamentally influence brain growth and function. Although this NPC diversity is critical for neocortex expansion, the molecular basis of NPC fate determination and diversity in the developing human brain remains a mystery.

Primary cilia extend from the surface of most vertebrate cells, originating from the mother centriole, which acts as a basal body that docks to the plasma membrane^17^. The docking process is mediated by a specialized, pinwheel-like structure called the distal appendages, formed by a group of distal appendage proteins (DAPs) at the distal end of the mother centriole^18,19^. DAPs are essential for initiating ciliogenesis, and their molecular composition and biogenesis mechanisms are only beginning to emerge ^20–22^. Primary cilia mediate several growth signaling pathways, including Sonic Hedgehog (SHH), which is vital for brain development^23,24^. Mutations in cilia-specific genes lead to ciliopathies, which are characterized by diverse phenotypes affecting multiple organs and are often accompanied by neurodevelopmental disorders ^25,26^. Studies have shown that specific ciliary dysfunctions are linked to neurodevelopmental disorders, known as "non-classical ciliopathies," emphasizing the vital role of cilia in human brain development ^17,27–29^. For instance, SHH signaling is crucial for controlling dorsal and ventral patterning during telencephalon development, and disruptions in this pathway can impair neural tube closure ^30–32^. Mutations in SCLT1, a component of DAPs, is associated with neuroretinal degeneration^28^. Importantly, many genes related to basal body-cilia are enriched in the developing brain, and their regulated dynamics contribute to drive the lateral growth of the cerebral cortex ^33,34^. This suggests that ciliary functions tailored to specific contexts, i.e., cell type-specific functions, may have evolved specifically to support brain development. This led us to search for unknown molecules that regulate ciliogenesis in the developing human brain.

In this study, we identify a previously uncharacterized gene, KIAA0408, and demonstrate that it encodes a 79-kilodalton (kDa) protein essential for regulating cilium formation, which we refer to as CCB79. CCB79 is mainly expressed in intermediate progenitors and both excitatory and inhibitory neurons located in the subventricular (SV) and outer subventricular (oSV) regions of the developing brain. Molecularly, CCB79 is a centriolar distal appendage protein (DAP) that is dynamically regulated during the cell cycle, appearing only at the start of ciliogenesis. Using proximity labeling and mass spectrometry, we show that CCB79 forms a complex with other DAP components. CRISPR/CAS9-mediated deletion of CCB79 disrupts the DAP and hinders cilium formation. Functional studies in genome-engineered brain organoids, combined with comparative single-cell (scRNA) and whole-transcriptome analyses, revealed abnormal ciliogenesis and misregulated SHH signaling, leading to altered NPC diversity and defects in brain tissue organization. We find that intermediate and ventral progenitors are particularly vulnerable when CCB79 is absent. Notably, the human gene can rescue cilia defects, while the mouse ortholog cannot. This suggests that, although CCB79 is a vertebrate gene, the primate variant has rapidly evolved to regulate specific aspects of ciliogenesis that are crucial for cortical expansion in primates.

### Discovery of CCB79 as a controller of cilium biogenesis

To identify new regulators of NPC diversity, we analyzed single-cell RNA sequencing (scRNA-seq) data from our early-stage brain organoids, described in Figures 5-6^35–37^. Uniform Manifold Approximation and Projection (UMAP) revealed six significant cell clusters, including telencephalic neural progenitors, dorsal and ventral progenitors, and excitatory and inhibitory neurons **(Figure 1A)**. Since intermediate progenitors are relatively abundant in gyrencephalic species with a high neurogenic capacity for neocortex expansion ^13–15^, we calculated the differentially expressed genes in intermediate progenitor-containing clusters (C5 and C6 of SVZ and oSVZ region) compared to the telencephalic progenitor cluster (C2 of apical ventricular region) **(Figure 1B)**. We identified a group of genes that were significantly upregulated. We could assign known functions to all of these genes, except for the *KIAA0408* mRNA **(Supplementary Table 1).** Computing the normalized expression of *KIAA0408* across various human tissues revealed that its expression is enriched in fetal brain tissues, specifically in early neuronal cell types and at different stages of fetal development^38^ **(Figure S1A-C).** This is corroborated by scRNA data derived from our brain organoids **(Figure 1C)**. Notably, in our UMAP, the expression of *KIAA0408* coincides with an enriched expression of EOMES, INSM1, NKX2-2, GAD2, and SLC17A6, markers of intermediate progenitors and excitatory and inhibitory neurons emerging from dorsal and ventral progenitors, respectively **(Figure 1D and S1D)**. A pseudotime trajectory analysis confirms this, indicating that our brain organoids harbor cell lineages that follow sequential stages of brain development, including dorsal and ventral patterning events **(Figure 1D)**.

**Figure 1.**
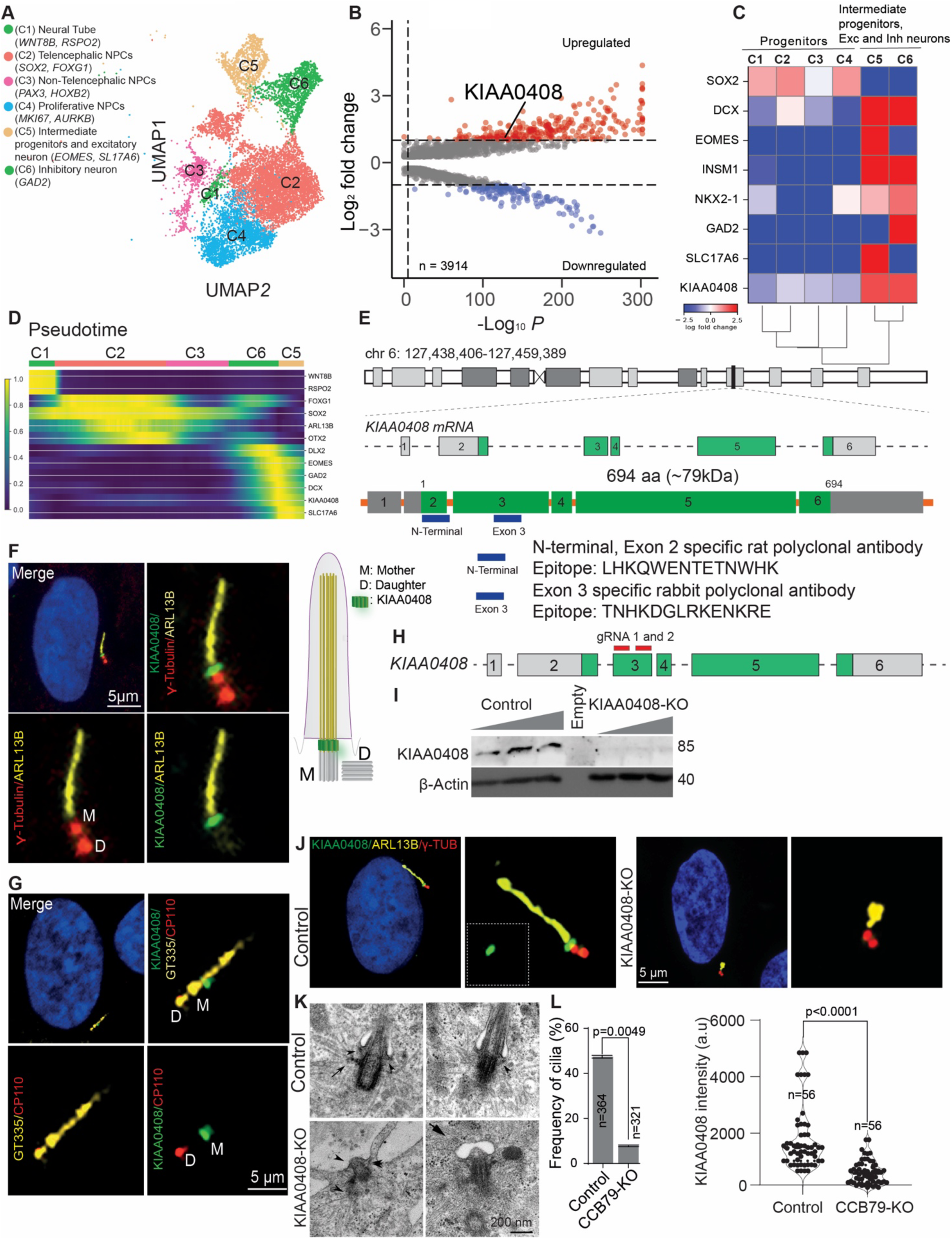
Discovery of CCB79 as a controller of cilium biogenesis. **A.** UMAP plot showing significant cell clusters derived from Day 14 human brain organoids. Cluster description and markers used to annotate the clusters are given in the legend. **B.** Volcano plot of differentially expressed genes between cluster 2 (telencephalic NPCs) and cluster 5 (Intermediate progenitors) identified *KIAA0408*, which has unknown functions. **C.** A matrix plot to display the relative expression levels of the considered genes. Note that *KIAA0408* is enriched in clusters 6 and 7, coinciding with intermediate progenitors, inhibitory, and excitatory neuronal markers. **D.** Pseudotime trajectory analysis of cell clusters showing cell lineages that follow sequential stages of brain development with dorsal and ventral patterning. **E.** Gene structure of *KIAA0408*. Predicted exons (green bar) are translated into a 79 kDa protein. The position and epitope sequences used to generate anti-KIAA0408 are given (blue line). **F.** Anti-KIAA0408 antibodies specifically recognize the endogenous protein located between the distal end of the mother centriole (γ-tubulin, red) and the ciliary base (ARL13B, yellow). Scale bar 5 µm. **G.** Anti-KIAA0408 antibodies specifically recognize the mother centriole, which is also labelled by GT335 (glutamylated tubulin, yellow). The daughter centriole is specified by CP110 (red), which is absent from the mother centriole. Scale bar, 5 µm. **H.** Position of guide RNAs used to ablate KIAA0408 using CRISPR/CAS9 is shown on its 3^rd^ exon structure. **I.** Anti-KIAA0408 generated against the C-terminal epitope (shown in E) recognizes a 79 kDa band only in unedited iPSCs, not in KIAA0408-KO iPSCs that lack the translated protein. **J.** KIAA0408-KO iPSCs lack KIAA0408 immunoreactivity (green in control) and a fully assembled cilium (Yellow in control). Mother and daughter centrioles are stained with γ-tubulin, shown in red; scale bar, 5 µm. **K.** Serial sectioning EM analysis reveals that control iPSCs (top panel) typically exhibit structurally assembled primary cilia with a mother centriole at their base. Arrowheads indicate ciliary pockets, and the arrow points to an appendage structure. KIAA0408-KO iPSCs (bottom panel) lack both appendage structures (arrows) and the cilium. Instead of a typical ciliary pocket, the mother centriole is docked to a membrane bulge. Scale bar, 200 nm. A representative image from at least 15 cells is displayed. **L.** Bar graphs display the frequencies of ciliated cells and KIAA0408 intensity between control and KIAA0408-KO cells. Data are shown as mean ± SEM from at least three independent experiments. For KIAA0408 intensities (scatter plot at right), a non-parametric Mann-Whitney test was used. The “n” numbers and p-values are provided in the graph.

KIAA0408 is an uncharacterized gene located on chromosome 6orf174, considered a complex locus because it merges with another gene, *MTCL3*. Although their coding regions do not overlap, a read-through RNA product—likely a non-coding RNA and a candidate for nonsense-mediated mRNA decay—is produced^39^. *KIAA0408* has six exons that encode a protein with coiled-coil domains **(Figure 1E)**. To determine its subcellular localization, we transiently expressed the human *KIAA0408* cDNA with a green fluorescent protein reporter tag at its N-terminus in human induced pluripotent stem cells (iPSCs). We observed the gene product as a distinct dot in each cell. To test whether these dots are basal bodies associated with primary cilia, we immunostained cilia using ARL13B antibodies and found that KIAA0408 localizes at the ciliary base **(Figure S1E)**. To label the endogenous protein, we generated polyclonal rat antibodies against the N-terminal peptide of KIAA0408 **(Figure 1E).** In these experiments, we used NPCs differentiated from iPSCs. Anti-KIAA0408 antibodies labeled KIAA0408 as a single, distinct dot at the distal end of the mother centriole, filling the space between the mother centriole (defined by γ-tubulin) and the cilium (defined by ARL13B) **(Figure 1F)**. We further immunostained the NPCs with antibodies against glutamylated tubulin GT335 and CP110. GT335 antibodies label the mother and daughter centrioles as well as the cilium. CP110, a centriolar protein, specifically labels the daughter centriole when the mother centriole serves as a template for cilium assembly^40,41^. This analysis reaffirmed the endogenous localization of KIAA0408, showing it labels the distal end of the mother centriole and connects it to the cilium **(Figure 1F-G).**

To validate the KIAA0408 antibody and determine the gene’s function, we knocked out the KIAA0408 gene in iPSCs using the CRISPR/CAS9 method using two guide RNAs targeting exon 3 ^42^ **(Figure 1H).** Isolation of pure clones and Sanger sequencing of the PCR-amplified region revealed that the gene editing resulted in a 35-nucleotide deletion between the two guide RNA binding sites in the targeted genomic region. We also observed the introduction of a stop codon within 22 bases of the deleted region, which stops translation after 90 amino acids **(Figure S2A)**. To detect the protein in Western blots, we raised polyclonal rabbit antibodies against the exon 3 peptides of KIAA0408 **(Figure 1E),** which recognized the protein bands only in unedited control iPSCs but not in knockout cells **(Figure 1I).** On the other hand, N-terminal antibodies did not show any immunoreactivity in KIAA0408-KO cells **(Figure 1J)**. If detected, there was only a faint signal, suggesting either reduced protein recruitment or that the residual signaling was due to the presence of the first 90 intact amino acids, which cover the epitope region for the antibody. As a result, unlike unedited cells, most KIAA0408-KO cells lacked primary cilia. When detected, the cilia were bulgy, short, thin, and long, indicating they are aberrant **(Figure 1J and S2B).** Ultrastructural investigation using transmission electron microscopy (EM) revealed that mother centrioles contained appendages in control cells that dock to the plasma membrane, form ciliary pockets, and emanate axonemal microtubules, and assemble cilia. In contrast, KIAA0408-KO cells lacked appendages in centrioles, forming a bulgy vesicle-like ciliary pocket, and failed to assemble structurally normal cilia **(Figure 1K)**. Finally, while the number of centrosomes remains unchanged in KIAA0408-KO cells, the frequency of ciliated cells and CCB79’s recruitment to mother centrioles is significantly reduced compared to controls **(Figure 1L and S2C).** These results indicate that KIAA0408 localizes to the distal end of the mother centrioles and is critical for initiating ciliogenesis. For clarity, we refer to KIAA0408 as CCB79, designating it as a **C**ontroller of **C**ilium **B**iogenesis with a molecular weight of 79 kDa.

### CCB79 is a cell cycle-regulated centriolar distal appendage protein

To gain mechanistic insight into how CCB79 initiates ciliogenesis, we used a proximity labeling technique ^43^ that combines biotinylation with mass spectrometry (MS) to map the spatial protein-protein interaction network of CCB79. We generated hTERT-immortalized retinal pigment epithelial cells (RPE-1) that express doxycycline-inducible miniTurbo, a biotin ligase fused to a FLAG-tagged CCB79. We purified the CCB79 proximity proteome 30 minutes after starting the biotinylation reaction in 4 biological replicates **(Figure 2A)**. After the MS detection, we performed enrichment analysis applying a p-value cutoff of 0.1 and Log_2_ fold change cutoff of 0.75 between miniTurbo control and miniTurbo-CCB79. Through this analysis, we found that CCB79 forms a complex with proteins crucial for cilium initiation, including distal centriolar and ciliary proteins, and notably, a significant proportion of centriolar appendage proteins **(Figure 2B-D and Supplementary Table 2)**. Because CCB79 localizes at the distal centriole **(Figures 1F and G)** and interacts with appendage and ciliary proteins, we hypothesized that CCB79 is an unrecognized component of the centriolar appendage, possibly involved in mediating a Cilium Assembly Complex (CAC) to initiate ciliogenesis. To map the precise localization of CCB79 within the mother centriole relative to centriolar appendage proteins, we used structured illumination microscopy (SIM) on NPCs. Compared to GT335 and CP110-specific staining (magenta), CCB79 (green) formed a distinct ring distal to the mother centriole at the base of the cilium **(Figure 2Ei-ii).** We then co-stained with ODF2 (magenta), which marks the subdistal appendages of the mother centriole ^21,44^. While the top view showed that the CCB79 ring is clearly distal to ODF2 signals, the tilted view revealed that OFD2 forms a hollow cylinder surrounded by the CCB79 ring **(Figure 2Eiii).** To further confirm CCB79’s position, we co-stained with CEP164 (magenta), a distal appendage protein. CEP164 formed a ring co-localizing with the CCB79 ring. Notably, the CCB79 ring displayed distinct periodic densities, suggesting they are distal appendages of the mother centriole **(Figure 2Eiv).** Finally, using super-resolution stimulated emission depletion (STED) microscopy, we visualized nine distinct densities of FBF1 (magenta), another distal appendage protein, which co-localized with CCB79 **(Figure 2Ev-vi).** Importantly, imaging several centrioles showed that the diameter of CCB79’s ring overlaps with the average diameter of CEP164 and FBF1 rings but not the OFD2 ring **(Figure 2F).** These results indicate that CCB79 is a component of the centriolar distal appendage structure at the base of the cilium **(Figure 2G-H).**

**Figure 2.**
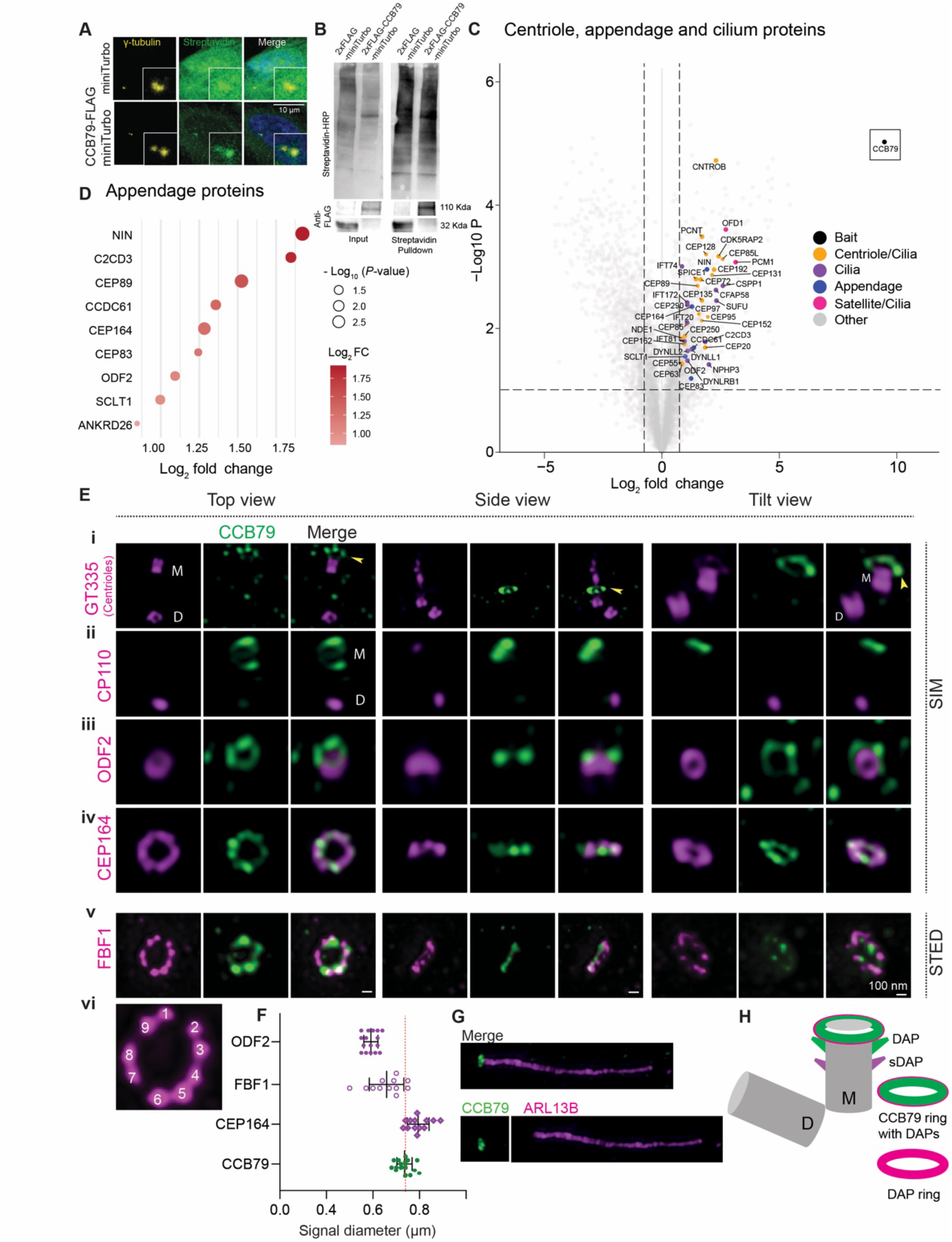
CCB79 is a centriolar DAP. **A.** RPE1 cells expressing miniTurbo (top) and CCB79-miniTurbo (bottom). Streptavidin labels the biotinylated proteins in cells (green). The biotinylation signal (green) localizes to the centrioles of cells expressing CCB79-FLAG-miniTurbo. Centrioles are marked by γ-tubulin (yellow). Insets show zoomed-in centrioles. Scale bar, 10 µm. **B.** Purification of CCB79’s proximity protein network. Cell extracts are shown to contain biotinylated proteins using streptavidin-HRP (Top blots). After affinity purification, anti-FLAG antibodies recognize CCB79-FLAG-miniTurbo (110kDA) (Bottom blots). Molecular weights are shown at right. **C.** Enrichment analysis following mass spectrometry. Volcano plots show proteins that are significantly enriched in complexes with CCB79. The bait, CCB79, is strongly enriched and highlighted with a box. Legends explain the different types of proteins. p value < 0.10 & Log_2_ Fold Change > 0.75. **D.** Centriolar appendage proteins and their fold change enrichments in the CCB79 proximity protein network. p value < 0.10 & Log_2_ Fold Change > 0.75. **E. (i)** SIM-based imaging of the mother (M) and daughter (D) centrioles specified by GT335 (magenta) and showing CCB79 (green) ring distal to the mother centriole, like a crown (Yellow arrowhead). Likewise, CP110 (magenta) labels only the daughter centriole that is distant from the CCB79 ring **(ii)**. OFD2, a subdistal appendage protein (magenta), is distinctly away from the CCB79 ring (green), forming a hollow tube projected into the CCB79 ring **(iii)**. CEP164, a distal appendage protein (magenta), forms a ring that co-localizes with the CCB79 ring (green). Both CEP164 and CC79 rings show periodic densities **(iv)**. STED-based imaging of FBF1, another centriolar DAP, forms a distinct nine-fold symmetrical structure of DAP that is co-localized with the CCB79 ring **(v-vi).** Top, side, and tilt views are shown for all examples. Scale bar 100 µm. **F.** The scatter plot shows the mean diameter of the CCB79 ring from each centriole imaged, which is similar to DAP molecules and wider than OFD2, a subdistal appendage protein (sDAP). **G.** A SIM image of CCB79 (green) and cilium (ARL13B, magenta) **H.** Schematic diagram showing positions of DAP (magenta), CCB79 (green), and sDAP rings at a mother centriole.

The primary cilium is a dynamic organelle that assembles and disassembles, closely linked to the cell cycle^45,46^. It assembles when the cell exits mitosis and disassembles upon re-entry into the cycle. Our expanded analysis of CCB79’s localization across different cell cycle stages revealed a single CCB79 dot in the G0/G1 phase, when cells build a cilium. Cells in G2, anaphase, and telophase lacked CCB79. Interestingly, CCB79 reappeared during late cytokinesis as a single dot in one of the dividing cells that began to form a cilium **(Figure S2D).** This pattern shows that the mother centriole is preparing to assemble a cilium in G1 by recruiting CCB79.

Appendage proteins specify mother centrioles competent to template cilium biogenesis ^19,20^. However, most appendage proteins not only localize to the mother centrioles at the onset of ciliogenesis but are present throughout the cell cycle. Comparing the localization behaviors of selected appendage proteins, such as CEP164 and FBF1, to CCB79 revealed that CCB79 intensity peaks at G0/G1, the stage when cells assemble cilia, and rapidly declines as cells proceed to S, G2, and M phases, where cells disassemble cilia **(Figures S2E and S3A)**.

To further confirm that CCB79 appears only at the beginning of cilium assembly, we synchronized RPE-1 cells by serum starvation to reach G0 and then stimulated them with serum to transition to G1-S. After 72 hours of serum starvation, cells assembled cilia with CCB79 prominently localized at the distal end of the mother centriole as a single dot. During subsequent hours of serum addition, CCB79 staining diminished, indicating cilium disassembly. CP110 is localized solely at the daughter centrioles at similar levels **(Figure S3B)**. We also examined CPAP, a direct interactor of CP110, in these assays. CPAP is a cilium disassembly factor whose levels gradually increased as cells disassembled the cilium^29,47^ **(Figure S3C-D)**. These experiments show that CCB79, a cilium initiation factor, appears mainly in the distal appendage at the onset of cilium assembly.

### CCB79 is a primate-specific cilium initiation factor

Since centriolar appendage proteins are essential regulators of cilium biogenesis, we conducted a phylogenetic analysis of centriolar DAPs with CCB79 to examine their conservation across different species. To achieve this, we employed clustering analysis to infer evolutionary models and identify gene expression patterns across various species^48^. The results showed that the CCB79 gene first appeared in frogs and then continued to evolve in higher vertebrates and terrestrial animals. Interestingly, the emergence of CCB79 expression coincided with other known core DAP molecules such as TTBK2, CEP164, FBF1, CEP83, CEP89, C2CD3, and SCLT1 **(Figure 3A).** This finding prompted us to analyze its amino acid sequence. The basic pairwise alignment of sequences from different species relative to humans revealed significant changes in the CCB79 protein during evolution. For example, a mouse protein, which shares 55% identity with humans, rapidly evolved into a marmoset protein with 94% identity to the human protein within approximately 48 million years. Evolutionary pressure then slowed from marmosets to humans through macaques, bonobos, and chimpanzees over the past 43 million years **(Figure 3A)**. Additionally, multiple sequence alignment of CCB79 across various species uncovered notable insertions, substitutions, and deletions in their amino acid sequences. Notably, primates, starting with Old World monkeys, gained a 43-amino-acid region in exon 5, which is absent in rodents **(Figure S4A).**

**Figure 3.**
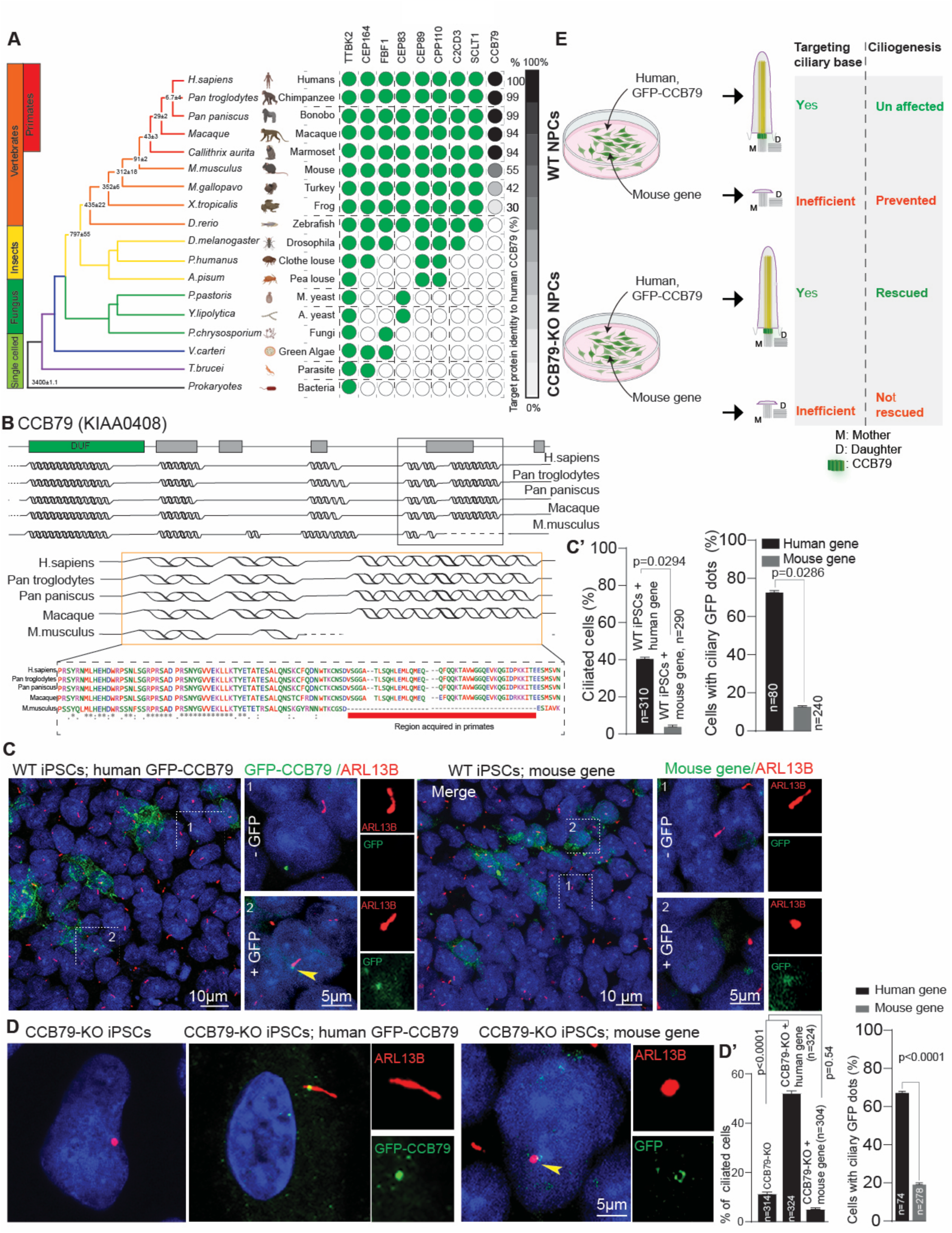
CCB79 is a vertebrate protein and a primate-specific cilium initiation factor. **A.** Phylogenetic conservation of CCB79 and other centriolar DAPs from bacteria, single-celled organisms, invertebrates, vertebrates, mammals, to humans, along with the known timeline of evolution. The last column shows the target protein (CCB79 from other species) and its percentage identity to the human CCB79 protein (represented by gradients of grey to black). Note, CCB79 appears only in amphibians and has evolved prominently in terrestrial animals. **B.** AlphaFold prediction of secondary structure of CCB79 between rodents and primates. Compared to rodents, primate genes gain 43 amino acids, which enable the formation of an alpha helix in primates. **C.** Inducible expression of GFP-CCB79 in wild-type iPSCs targets the ciliary base as a single dot (yellow arrowhead). The mouse gene (right panel) does not target the ciliary base efficiently and abolishes ciliogenesis. Both transfected and untransfected cells are shown, with the scale bar indicated in the panels (10 and 5 µm). Results are quantified in **C’**. Data presented as mean ± SEM from at least three independent experiments. For the frequencies of ciliated cells, a non-parametric T-test was used; for cells with ciliary GFP-CCB79 dots, a non-parametric Mann-Whitney test was used. The “n” numbers and p-values are provided in the graph. **D.** Human CCB79 expression in CCB79-KO rescues the cilia loss by targeting the mother centrioles and cilium initiation. The mouse gene (right panel) does not rescue the loss of cilia. Results are quantified in **D’** at right. Data presented as mean ± SEM from at least three independent experiments. For the frequencies of ciliated cells and cells with ciliary GFP-CCB79 dots, a non-parametric Mann-Whitney test was performed. The “n” numbers and p-values are provided in the graph.

We therefore focused our analysis on exon 5. Alignment of the human CCB79 gene with its mouse counterpart shows that the human gene has gained an extension of 131 nucleotides at its C-terminus **(Figure S4B, green extension)**. This suggests that the transcript may have arisen due to small variations at the splice sites caused by point mutations. Analyzing the potential splice site revealed that there is a donor splice site in exon 5 of the mouse gene (we call this mSD5), which is suppressed in humans by a mutation—specifically, a nucleotide exchange (+3A>C), a point mutation three nucleotides downstream of the 5’ splice site **(Figure S4B).** This is further supported by a decrease in the Hbond score (HBS), which measures complementarity to U1 snRNA of the spliceosome complex. Alternatively, this suppressed site is compensated by a novel splice site gained through another mutation further downstream at position +1 (SD5: +1T>G) in the human gene. This results in the exonization of intron 5 in the mouse transcript. HEXplorer plots in Figure S3B highlight the changes in the splicing environment caused by the mutations in human SD5 and mSD5. In summary, compared to the mouse gene, the human CCB79 gained 131 nucleotides at position 127,446,277 in the 5^th^ exon, located on chromosome 6, leading to a new exonization and the addition of 43 amino acids to the translated protein.

To predict how the gain of 43 aa could alter the structure and folding characterization of the protein in primates, we utilized the AlphaFold protein structure prediction tool. This analysis revealed a gain of alpha-helices in primates, indicating that this alpha-helix can be functionally traced back to a common ancestor of Old-World monkeys but is absent in rodents **(Figure 3B).** Therefore, with the gain of a 43 aa containing alpha-helix, it is plausible that the CCB79 is a primate-specific transcript essential for ciliogenesis in primates and is functionally different from the rodent gene. Thus, although the CCB79 is a vertebrate gene, the human or primate transcript has undergone significant evolutionary pressure to acquire primate-specific functions **(Figure S4C-D)**.

To test this experimentally, we cloned the mouse ortholog and examined whether it could rescue the ciliogenesis defects of CCB79-KO iPSCs. First, we conducted our experiments in wild-type iPSCs. While the human gene CCB79 effectively targeted the ciliary base, the mouse gene did not do so efficiently. Furthermore, the mouse gene inhibited the cells from forming cilia **(Figure 3C)**. Accordingly, while the human CCB79 fully restored ciliogenesis in CCB79-KO iPSCs, the mouse gene neither targeted the ciliary base effectively nor rescued the loss of cilia. These experiments reveal functional differences between human and mouse transcripts, and CCB79 is a primate-specific factor involved in cilium initiation **(Figure 3D).** Figure 3E summarizes the findings. Obtaining stable clones of iPSCs expressing CCB79 was impossible, as overexpression did not produce healthy lines. Therefore, we expressed the gene in a doxycycline-inducible manner and analyzed a mixed population of cells.

### CCB79 removal is sufficient to trigger NPCs to undergo premature differentiation and abolish cilia initiation

We conducted a series of experiments to characterize the loss of function of CCB79 and its role in cilium initiation and NPC maintenance using iPSC-derived NPCs. While both variants (control and CCB79-KO) did not show a significant difference in EdU incorporation, unlike controls, CCB79-KO NPCs spontaneously differentiated into neurons at the expense of self-renewing NPCs. Notably, these experiments were performed under conditions that do not promote differentiation, indicating that CCB79 is essential for maintaining NPCs in their self-renewal state **(Figure 4A).**

**Figure 4.**
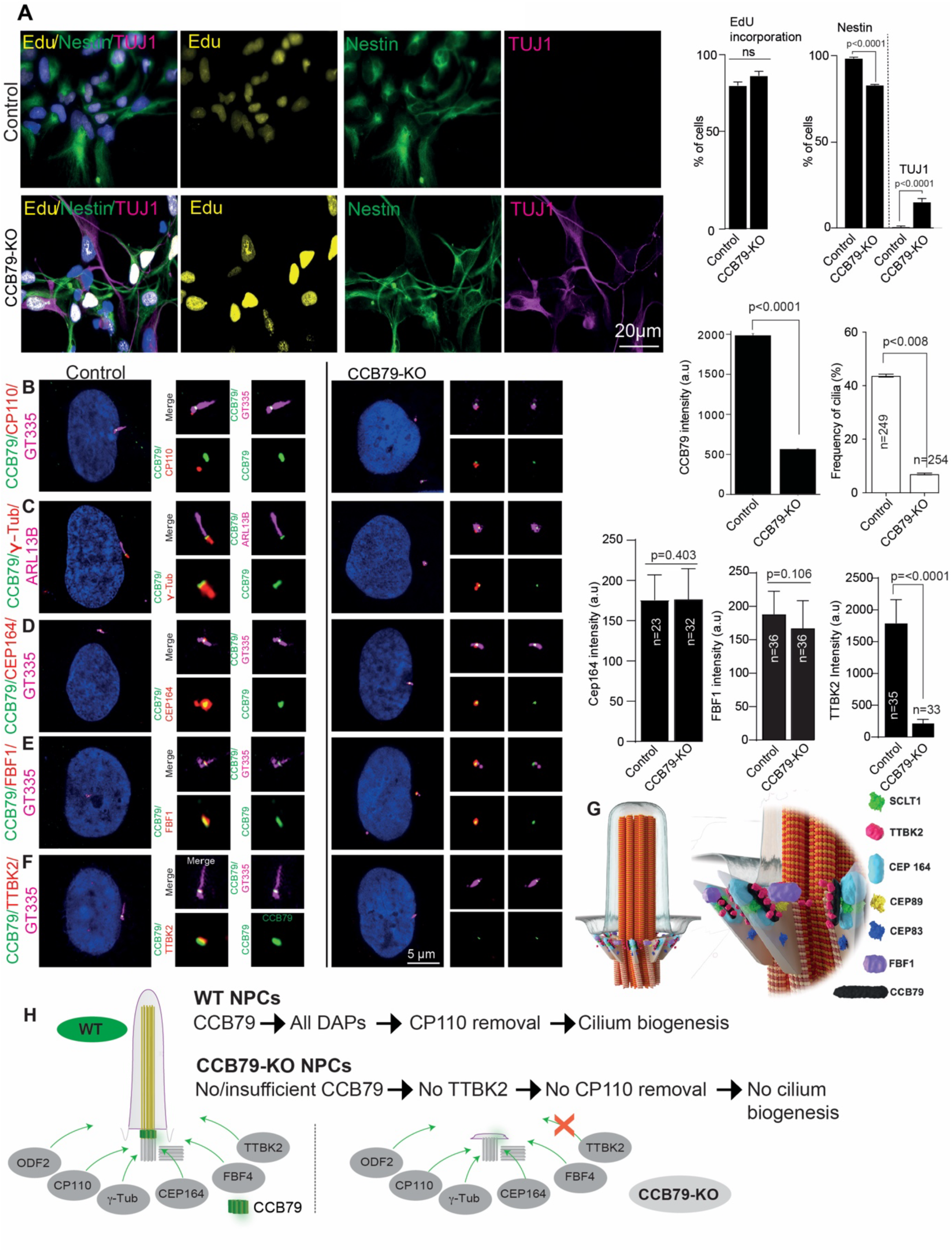
CCB79 maintains NPCs self-renewal and initiates ciliogenesis. **A.** CCB79 is critical for maintaining NPCs in their self-renewal state. Under non-differentiating conditions, control NPCs self-renew. CCB79-KO NPCs undergo spontaneous differentiation into neurons (TUJ1, magenta), leading to the loss of NPCs (Nestin, green). Edu (yellow) incorporation does not differ between control and CCB79-KO NPCs. Bar graphs at right quantify the findings. Data presented as mean ± SEM from at least three independent experiments. For quantifying Nestin versus TUJ-1 positive cells, a non-parametric Mann-Whitney test was performed. The “n” numbers and p-values are provided in the graph. **B-F.** DAP recruitment assay. Control NPCs recruit CP110, a centriolar protein (red, **B),** γ-tubulin (red, **C),** CEP164 (red, **D),** FBF1 (red, **E)**, and TTBK2 (red, **F)** (red) to the base of the cilia (ARL13B or GT335, magenta). CP110 labels only the daughter centrioles when the mother centriole templates ciliogenesis. In all cases, CCB79 is recruited to the mother centrioles at the ciliary base (green). The right panels show CCB79-KO NPCs, which either fail or faintly recruit CCB79 (green), and cells have either no cilia or aberrant cilia (magenta). Except for TTBK2 (red, **F),** the recruitment of other components is not significantly perturbed. Note that CP110 remains persistent in the mother centrioles that do not assemble cilia. Scale bar 5 µm. Bar diagrams on the right quantify the findings. Data presented as mean ± SEM from at least three independent experiments in each case. For intensity measurements (CCB79, CEP164, FBF1, and TTBK2), a non-parametric Mann-Whitney test was performed. The “n” numbers and p-values are provided in the graph. **G.** The drawing shows a mother centriole templating a cilium with the hypothetical topology of various DAP components, as indicated by CCB79. Legends are provided to the right to aid in understanding the experimental findings. **H.** The schematic summarizes the experimental results where CCB79-KO NPCs specifically fail to recruit TTBK2, and thus, no cilium is initiated.

Next, we aimed to dissect the underlying mechanisms of defective ciliogenesis. To achieve this, we conducted DAP recruitment assays in cells in the presence and absence of CCB79 since CCB79 appears to form a cilium assembly complex (CAC) with other DAP proteins **(Figure 2A-D).** CCB79-KO NPCs revealed the following findings in comparison to healthy controls **(Figure 4B-F)**. ***i)*** Impaired CCB79 recruitment, meaning CCB79-KO NPCs recruited either no or significantly reduced levels of CCB79 (green, CCB79, panels B-F); ***ii)*** Aberrant cilia, wherein cilia were either absent or presented as a dot or bulging short structure (magenta, GT335 or ARL13B, panels B-F); ***iii)*** CP110 was never removed but remained on both daughter and mother centrioles (red, CP110, panel B); ***iv)*** Recruitment of CEP164 and FBF1 were affected but not significantly (red, panels C, D, and E); and ***v)*** Strikingly, CCB79-KO NPCs failed to recruit TTBK2 (red, panel F). Panels on the right quantify the findings. In summary, these results indicate that CCB79 is a critical factor in cilium initiation, and its loss of function specifically inhibits TTBK2 recruitment, thereby impairing ciliogenesis. TTBK2 is an essential kinase that orchestrates cilia growth, and its absence abolishes cilium initiation^31^. Panel G presents a hypothetical topology of CCB79 as part of the DAP complex, initiating a cilium, and Panel H summarizes the findings.

### CCB79 removal downregulates telencephalic progenitor genes, ciliopathy genes, and perturbs Hedgehog signaling in 3D human brain organoids

To gain mechanistic insights into how CCB79 can maintain NPC diversity in 3D brain tissues, we employed unbiased mRNA transcriptomics at both bulk and single-cell levels (scRNA) on 14-day-old brain organoids cultured for 14 days in spinner flasks (see Methods). We generated these organoids from control and CCB79-KO iPSCs using our previously described method ^35–37^, as detailed in Figure 6. Briefly, we directly induced the differentiation of iPSCs into neural epithelium. Early-stage brain organoids generated using this method expressed a diverse range of cell types, including telencephalic neural progenitors, as well as dorsal and ventral progenitors, and excitatory and inhibitory neurons. These organoids mirror the sequential stages of telencephalon development, exhibiting both dorsal and ventral patterning events, and could model developmental disorders such as microcephaly **(Figure 1A, C-D)**.

The CCB79-KO organoids showed less growth compared to age-matched healthy controls, indicating developmental issues **(Figure S5A).** Thin sectioning and immunostaining for SOX2-positive progenitors, TUJ1-positive neurons, and ARL13-positive primary cilia revealed that control organoids had well-organized, expanded ventricular zones (VZ) rich in SOX2-positive progenitors, ARL13B-primary cilia in their apical lumen of the VZ, and a thin layer of TUJ-positive primitive cortical plate at the basal side, mirroring the cytoarchitecture of developing brain tissue. In contrast, the CCB79-KO organoids showed disrupted cytoarchitecture, unrecognizable VZs, and barely visible cilia in their lumen, with a noticeable reduction in SOX2 intensities but an increase in TUJ1-positive neurons throughout the tissue, indicating early progenitor differentiation and resulting NPC loss **(Figure S5B).**

To examine the molecular changes caused by CCB79 removal, we conducted a differential gene expression (DGE) analysis comparing control and CCB79-KO brain organoids. We identified a down-regulation of pan-telencephalic genes (FOXG1), dorsal telencephalic progenitor genes specifying newly born cortical projection neurons (NEUROD6), ventral telencephalon progenitor genes (GSX2 and DLX family) crucial for the development of forebrain GABAergic interneurons, genes relevant to intermediate progenitors and early-born cortical (TBR1), inhibitory (GAD2 and GABRA1), excitatory (SLC6A17), and inter (Calbindin) neurons. Gene ontology ranking aligned with the DEG analysis **(Figure S5C).**

We also observed a decrease in the expression of known ciliopathy genes, which supports the finding that CCB79 loss prevents cilium formation **(Figure S5D).** Interestingly, GPR161 is among the most downregulated genes. GPR161 functions as a negative regulator of Sonic Hedgehog (SHH) signaling, and its loss in mice results in increased SHH activity in the neural tube^49,50^. Similarly, increased SHH signaling in cortical organoids caused by the loss of INPP5E, a ciliary gene, results in ventralization^51^. Therefore, we examined SHH target genes (PTCH1 and GLI1), ligands (DHH, SHH, and IHH), and receptors (PTCH1 and SMO), and found their levels elevated, indicating that SHH signaling is disrupted in organoid tissues after CCB79 removal **(Figure S5E).** To confirm the increase in SHH components and activity, we calculated a weighted SHH pathway activation score and combined it with the normalized expression of canonical SHH pathway genes^52^. This analysis showed a significant increase in SHH activity in CCB79-KO organoids **(Figure S5F)**, driven by higher levels of key SHH signaling molecules. Notably, SHH pathway scores were inversely related to the expression of forebrain patterning transcription factors, including FOXG1, LHX6, and TBR1, indicating a coordinated disruption of SHH signaling and neurodevelopmental programs, consistent with the known role of primary cilia in brain development **(Figure S5G)**^27,31,50,51^. Collectively, these findings suggest that CCB79 loss causes de-repression of the SHH pathway and impairs forebrain gene expression, aligning with previously described ciliopathy phenotypes^53^. These changes in gene expression and ectopic SHH signaling could influence cell diversity in 3D brain tissue.

### CCB79 removal derails progenitor fate determination, disrupts developmental patterning, and changes cell diversity in 3D human brain organoids

To analyze changes in cell diversity, we examined the scRNA transcriptomes of approximately 18,000 control and 8,900 CCB79-KO cells isolated from 14-day-old brain organoids, tracking developmental changes caused by the removal of CCB79. We assigned the sequenced cells to neural tube (WNT8B, RSPO2), neural crest (PAX3, COLEC12), progenitors of telencephalic (SOX2, FOXG1) and non-telencephalic regions (PAX3, HOXB2), proliferative (MKI67, AURKB), and immature neurons from both telencephalic (STMN, DLX5, GAD1) and non-telencephalic origins (STMN2, LHX1) **(Figure 5A and S6-S7).** By evaluating similarities and differences, we found abnormal levels of oligodendrocyte precursors and neural crest lineages (clusters C2 and C9), with a simultaneous reduction in telencephalic progenitors and immature neurons derived from ventral progenitors (clusters C3 and C6). The unusual appearance of oligodendrocyte precursors may be linked to increased SHH signaling in ventral progenitors, which is known to induce premature specification of oligodendrocyte precursors at the expense of inhibitory neurons. This aligns with the effects of cilia-mediated SHH enhancement observed in cortical organoids and the chick ventral spinal cord^51,54^. We then quantified the differential expression of SHH genes across our cell clusters and, notably, identified elevated levels of SHH signaling components in specific cell types within cluster 9, which contains oligodendrocyte progenitors, in CCB79-KO organoids (**Figure S8)**.

**Figure 5.**
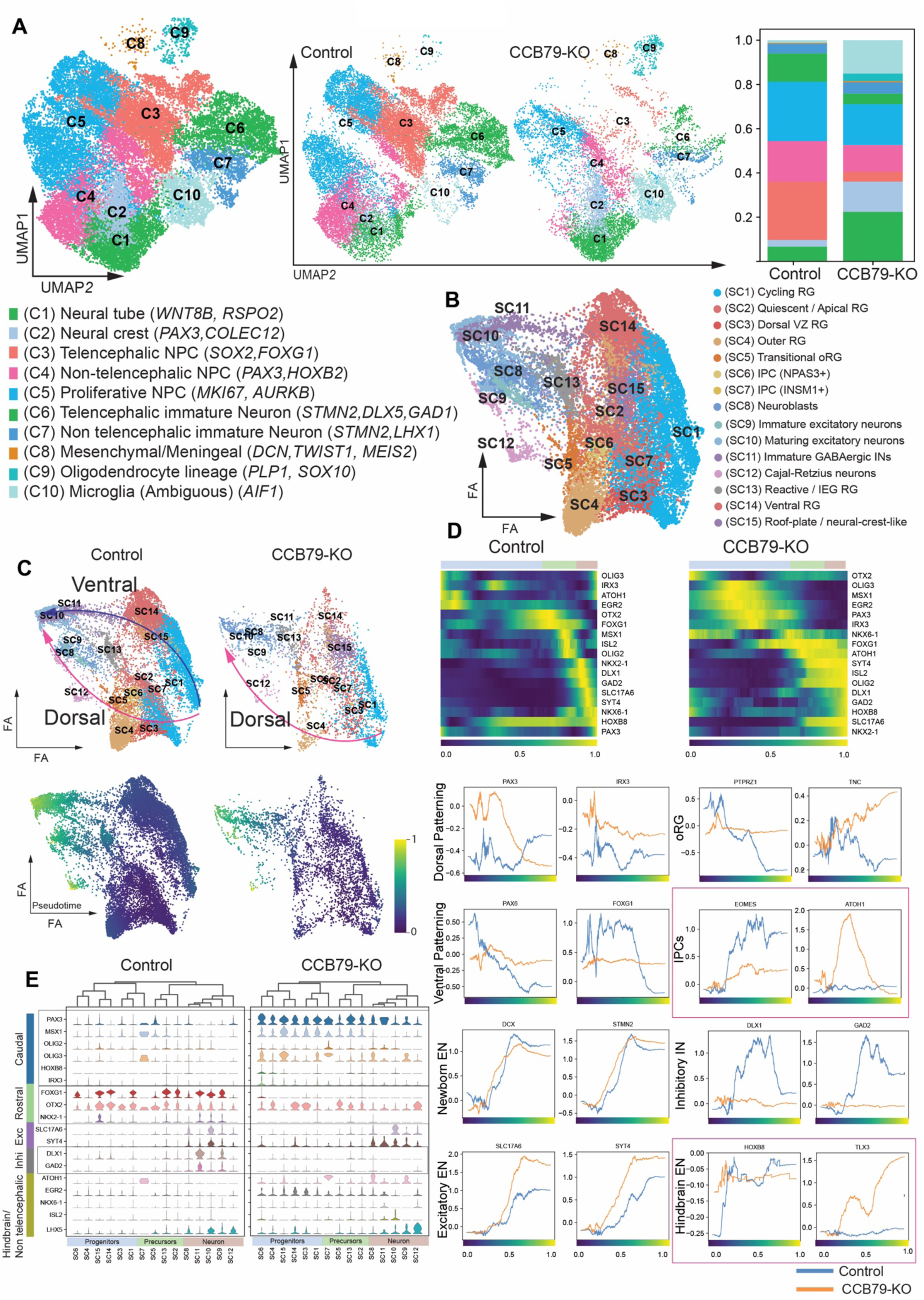
CCB79 is crucial for maintaining telencephalic progenitors and ventral patterning. **A.** Merged UMAP displaying various cell clusters. The legends below show the annotated cell clusters with the corresponding markers used. UMAP cell clusters of control and CCB-KO (middle panels) and proportions of various cell types between control and CCB79-KO organoids (right). **B.** Sub-iteration analysis of clusters C3, C4, C5, C6, and C7 identifies 15 subclusters. The legends at right show the annotated cell clusters. **C.** Pseudotime computation reveals regulatory trajectories exhibiting dorsal (magenta line) and ventral (blue line) patterning in the control. CCB79-KO shows a perturbed ventral patterning. **D.** Pseudotime plot and line plots of pseudotime expression trends for individual markers using a rolling mean (window size = 500 cells), faceted by gene and colored by condition (control = blue, CCB79-KO = orange). **E.** Stacked violin plots displaying expression distributions of the marker across annotated cell types in control (left) and CCB79-KO (right). Markers highlighting categories: caudal patterning (steel blue), rostral TFs (light green), excitatory neurons (purple), inhibitory neurons (grey), and hindbrain EN (olive green). Interesting regions of discussion are boxed.

Since CCB79 is enriched in SVZ and oSVZ progenitors, as well as excitatory and inhibitory neurons derived from both dorsal and ventral progenitors, we aimed to precisely map the changes in cell diversity in these regions. First, we performed a detailed analysis of clusters C3, C4, C5, C6, and C7 **(Figures 5B and S9)**. Next, we calculated pseudotime to project all cortical lineage cells onto a continuous differentiation axis that begins from the ventricular zone (VZ), extends through the inner and outer subventricular zones (SVZ/oSVZ), and ends in postmitotic neurons. Control organoids follow a regulated trajectory with dorsal and ventral patterning, reflecting the typical sequence of corticogenesis seen in human organoids and fetal cortex^55,56^. The first trajectory (blue line), which, which express markers of ventral radial glia, leading to the formation of GABAergic inhibitory neurons. The second trajectory (magenta line) indicates markers of dorsal and outer radial glia, which lead to excitatory neurons. Importantly, ventral patterning is disrupted in CCB79-KO organoids **(Figures 5C-D).**

In analyzing dorsal patterning events, we observed a clear, genotype-specific shift in the transcription factors of intermediate progenitors (IPs). In control samples, EOMES is a key transcription factor of IPs that promotes the specification of upper-layer excitatory neurons during forebrain development. In contrast, CCB79-KO cells expressed low levels of EOMES but an elevated level of ATOH1 along with its downstream regulator TLX3. ATOH1 is a transcription factor usually restricted to hindbrain neurogenesis, which enforces glutamatergic identity in dorsal spinal and hindbrain neurons **(Boxed line graphs in Figure 5D)** ^57–59^. This signature suggests that the loss of CCB79 also disrupts dorsally specified IPs but induces an ATOH1-driven progenitor program, which interferes with the typical EOMES-driven expansion phase in the SVZ/oSVZ. This imbalance likely causes shifts in dorsal cell types by reducing the number of classical IPs and upper-layer neurons while increasing atypical progenitors that are stalled in dorsal identity.

Furthermore, measuring the relative expressions of identity markers in progenitors, precursors, and neurons showed that control organoids exhibit rostral patterning events (identified by FOXG1 and OTX2). In contrast, CCB79-KO organoids shifted from rostral to caudal patterning, accompanied by an increase in hindbrain progenitors (HOXB1 and MSX1) and a decrease in inhibitory interneurons (DLX1 and GAD2) **(Figure 5E)**. We suspected that the observed misdifferentiation could cause defects in the cytoarchitectural organization of the CCB79-KO organoids.

To map this, we calculated the relative proportion of cell types in specific brain regions during development **(Figure S10A)**. We observed a significant depletion of cells in the VZ of CCB79-KO organoids, along with a downregulation of early progenitor markers, including PAX6 and FOXG1. This suggests either premature differentiation or impaired maintenance of radial glial cells, leading to a reduced progenitor pool **(Figure S10B)**. The increased presence of cells in the intermediate zone (IZ) and migratory zone (MZ) coincides with a sharp decline in the tangential migratory stream, suggesting that migration has stalled. Differential gene expression analysis reveals a downregulation of key neurogenic transcription factors (SATB2, DCX, DLX1/2) and neuronal markers (NEUROD1/2), as well as interneuron-specific transcription factors (LHX6, ARX), indicating delayed or impaired neuronal migration to the cortical plate **(Figure S10C)**. Additionally, marker-based scoring of division modes of radial glial cells showed an increase in asymmetric divisions and a decrease in symmetric divisions in CCB79-KO, indicating that neuronal commitment expands at the expense of self-renewing progenitors **(Figure S10D).** Overall, these findings demonstrate that CCB79 deletion disrupts the neurogenesis program through combined defects in progenitor dynamics, transcriptional regulation, and regional signaling, emphasizing the essential role of primary cilia in coordinating human cortical development. The schematic summarizes potential events that occur when CCB79 is removed **(Figure S11).**

### CCB79 ablation perturbs brain tissue organization due to NPC depletion, and the mouse ortholog does not rescue the defects

To verify the findings from our transcriptomic data and to characterize the effects of impaired ciliogenesis on 3D tissues, we performed imaging-based analyses of cell composition and cytoarchitecture. To visualize the entire brain organoid’s cytoarchitecture in its intact form, we tissue-cleared them, immunostained with SOX2 (progenitors), ARL13B (cilia), and MAP2 (neurons), and imaged with a light-sheet microscope. Despite its low resolution, we were able to volumetrically visualize the cytoarchitecture and measure the VZ, including its diameter and lumen size. Compared to controls, CCB79-KO organoids showed disrupted organization, characterized by fewer but thinner rosettes and an irregular distribution of neurons **(Figure 6A-D and movies 1-2)**. Since the primate CCB79 has undergone significant changes compared to its rodent ortholog, we tested whether the rodent gene could functionally rescue brain organization defects in 3D tissues. For this, we generated brain organoids using CCB79-KO iPSCs that express human CCB79 and mouse genes, as shown in Figure 3D. As stably expressing clones were not obtainable, we engineered them to express the genes in an inducible manner using doxycycline. While the human gene significantly rescued the rosette defects, it did not rescue defects related to the VZ. Conversely, the mouse gene failed to rescue any of the defects and even worsened the phenotype of CCB79-KO organoids, reaffirming our results with 2D cells described in Figure 3D **(Figure 6E-F).**

**Figure 6.**
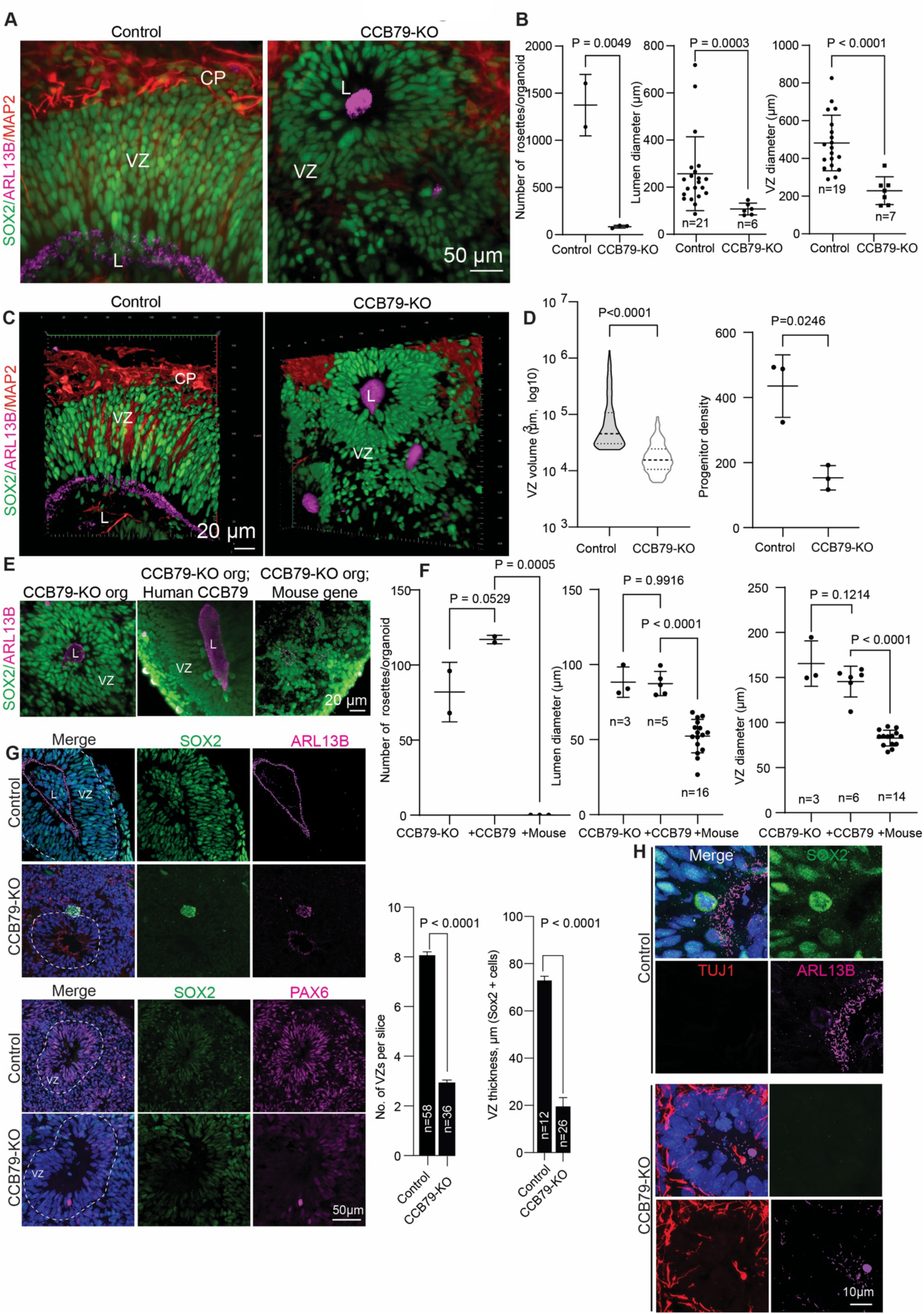
CCB79 ablation perturbs brain tissue organization, and the mouse gene does not rescue the defects. **A.** Tissue clearing and light sheet imaging of control and CCB79-KO brain organoids. Control organoids show defined VZs of several cell layers thick with a defined lumen at the apical side (SOX2, green). The VZ lumen (L) is defined by primary cilia (ARL13B, magenta). MAP2 defined the primitive cortical plate (CP) at the basal side (red). In contrast, CCB79-KO organoids do not exhibit a defined VZ, but rather rosette-like structures, which are significantly fewer in number than in the control: scale bar, 50 µm. **B.** Scattered plots quantify the number and dimensions of VZ between control and CCB79-KO. Data presented as mean ± SEM from at least three independent experiments. An unpaired Student’s t-test was used. The “n” numbers and the “p” values are given in the graph. **C.** Volume rendered images of panel A in a 3D representation. Refer to the corresponding movies, 1 and 2. **D.** Quantification of VZ volume and progenitor density between control and CCB79-KO after volume rendering. Data presented as mean ± SEM from at least three independent experiments. An unpaired Student’s t-test was used. The “n” numbers and the “p” values are given in the graph. **E.** While the human CCB79 partially rescues the defects of CCB79-KO, the mouse gene fails to rescue these defects, including the number of neural rosettes, lumen diameter, and VZ diameter. Scale bar 20 µm. **F.** Scatter plots quantify the rescue effects. Data presented as mean ± SEM from at least three independent experiments. One-Way ANOVA with Multiple Comparisons test. The “n” numbers and the “p” values are given in the graph. **G.** Thin sectioning and immunostaining of control and CCB79-KO organoids. Control organoids show typical cytoarchitecture, but CCB79-KO organoids do not. The number of VZ and its thickness are reduced. Besides, both SOX2 (green) and PAX6 (magenta)-positive progenitors are weak in CCB79-KO sections. Scale bar 50 µm. Bar diagrams at right quantify the findings between control and CCB79-KO organoids. Data presented as mean ± SEM from at least three independent experiments. A nonparametric T-test, followed by a Mann-Whitney test, was performed. The “n” numbers and the “p” values are given in the graph. **H.** Zoomed-in images of VZ lumen show a well-defined VZ lumen whose apical region is lined by primary cilia (ARL13B, magenta) in control organoids (Upper panel). SOX2-positive (green) progenitors define VZ. In contrast, CCB79-KO organoids (Bottom panel) are not structurally normal, progenitors lack SOX2, lumens lack cilia, and TUJ1-positive neurons are all over. Scale bar 10 µm.

We then analyzed the distribution of PAX6, SOX2, ARL13B, and TUJ1-positive cells in organoid slices and quantified their numbers. We found that CCB79-KO organoids exhibit perturbed ciliogenesis characterized by barely visible cilia in their lumen. Additionally, CCB79-KO organoids exhibited a disrupted and thinner VZ, with only a few layers of radial glial cells, and a noticeable decrease in PAX6/SOX2 intensities **(Figure 6G-H**).

We then investigated premature differentiation of NPCs as suggested by our scRNA analysis **(F/igure 11D)**. We found that a TUJ1-positive region is confined to the basal side of control organoids, forming a primitive cortical plate (CP) separated from the VZ. In contrast, CCB79-KO organoids showed neurons throughout the organoids, including within the VZ, along with a decrease in radial glial progenitors **(Figure 7A).** Since CCB79 is enriched in intermediate progenitors, we examined the presence of these cells. In controls, TBR2-positive cells form a layer of the subventricular zone (SVZ) basal to apical progenitors lining the ventricular lumen. Conversely, CCB79-KO organoids did not display a TBR2-positive layer. Similarly, scoring for PTPRZ1-positive outer radial glial cells revealed a marked reduction in population in both mutant organoids **(Figure 7B-D).** Therefore, the finding of increased neuron levels combined with progenitor depletion indicates that an aberrant, indirect, and premature neurogenesis occurs in organoids lacking functional CCB79 protein.

**Figure 7.**
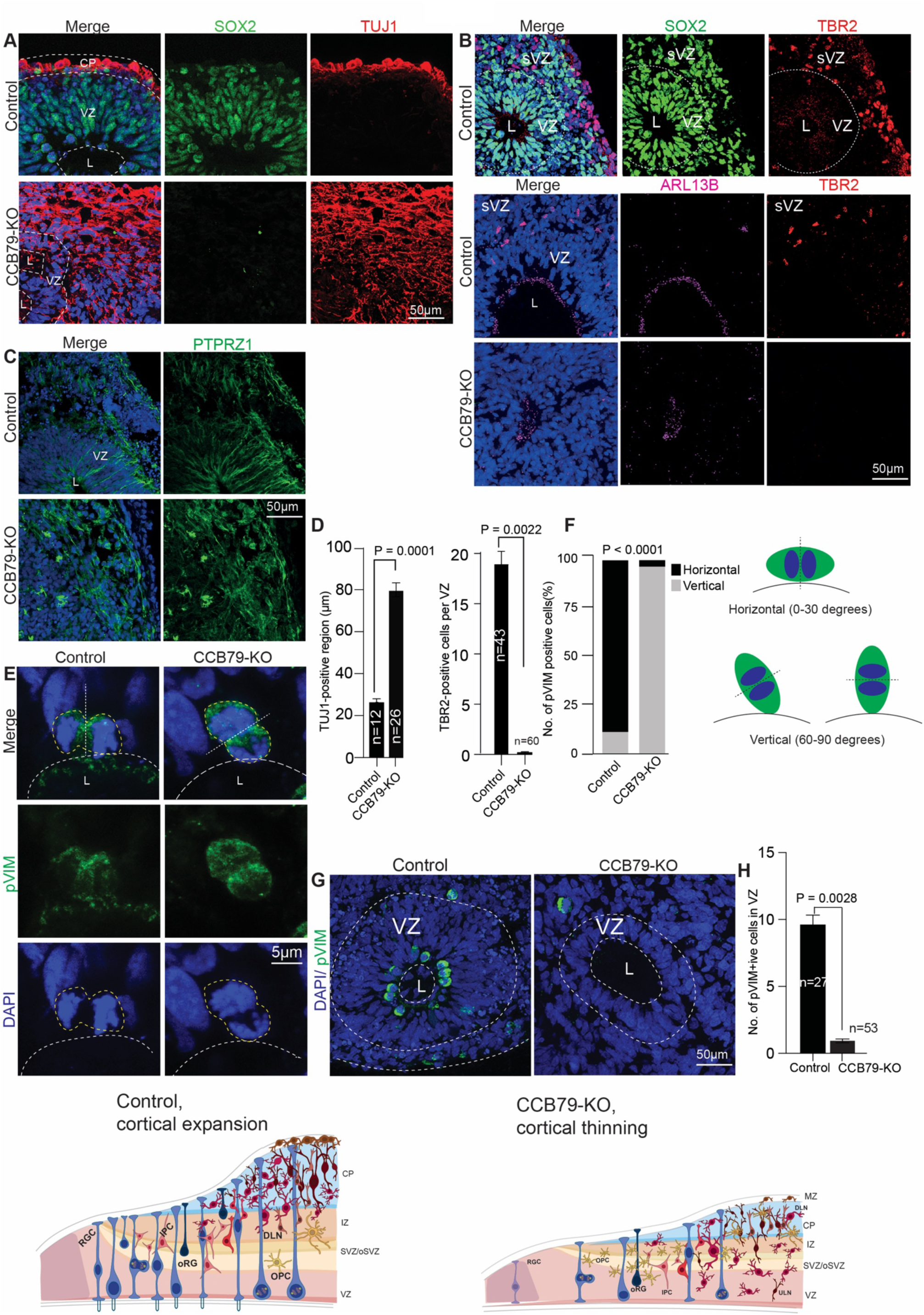
Progenitors in CCB79-KO brain organoids undergo premature differentiation. **A.** Control brain organoids show their typical cytoarchitecture with distinct VZ (SOX2, green) and TUJ1-positive (red) primitive cortical plate (CP). CCB79-KO organoids exhibit neurons throughout with a thinner VZ and a deficiency of SOX2-positive progenitors. Scale bar 50 µm. **B.** TBR2 (red) defines intermediate progenitors at the sVZ in controls. VZ is marked by SOX2-positive progenitors (green). Cilia (ARL13B, magenta) mark the VZ lumen. CCB79-KO organoids lack TBR2-positive cells. **C.** Control organoids display PTPRZ1, which is expressed in outer radial glial cells spanning from the VZ to the basal region (green). CCB79-KO organoids show a drastically reduced population of PTPRZ1 cells. **D.** Bar diagrams quantify the findings from panels A, B, and C. Data are presented as mean ± SEM from at least three independent experiments. A non-parametric Mann-Whitney test was performed. The “n” numbers and p-values are provided in the graph. **E.** Measuring premature differentiation by analyzing the kinetics of the apical progenitor division plane. Phospho-Vimentin (green) selectively labels the dividing apical progenitors at the VZ’s apical side. Control organoids predominantly display dividing progenitors whose division plane is horizontal to the VZ lumen (Left panel). CCB79-KO brain organoids (Right panel) show apical progenitors whose division plane is mainly vertical to the VZ lumen. Scale bar 5 µm. **F.** The bar diagram quantifies the distribution of the division plane. Horizontal and vertical. A schematic on the right shows horizontal and vertical division planes. Data are presented as proportion mean ± SEM from at least three independent experiments. Fisher’s exact test was performed. The “n” numbers and the “p” values are given in the graph. **G.** Control organoids, in general, have numerous Phospho-Vimentin-positive (green) apical progenitors, whereas CCB79-KO do not. Scale bar 50 µm. **H.** A bar diagram quantifies the distribution of Phospho-Vimentin-positive apical progenitors. Data are presented as mean ± SEM from at least three independent experiments. A non-parametric Mann-Whitney test was performed. The “n” numbers and p-values are provided in the graph. **I.** A schematic representation of the normal brain tissue development (Left) compared to the brain tissue in the absence of CCB79 (Right). This illustration should be viewed in conjunction with Figure S12 for a more detailed understanding of the defects caused by the loss of CCB79.

To explore the basis of premature differentiation, we studied the kinetics of apical progenitors whose division plane mostly orients horizontally toward the ventricular lumen. This orientation suggests a symmetric expansion of apical progenitors, leading to the formation of various NPC types critical for building a layered neocortex. Phospho-vimentin marks dividing apical progenitors, and by analyzing the division plane relative to the ventricular lumen, we determined the proportion of apical progenitors oriented horizontally (proliferating) versus vertically (differentiating) (Refs). While control organoids mainly exhibited horizontally oriented apical progenitors, CCB79-KO organoids contained cells oriented vertically, indicating a tendency to differentiate and be displaced from the apical side. This supports the finding that control organoids had significantly more Phospho-vimentin-positive cells than the CCB79-KO organoids **(Figure 7E-H)**. These results suggest that the loss of functional CCB79 impairs ciliogenesis, which subsequently triggers NPC differentiation, leading to NPC depletion and ultimately resulting in defective brain tissue organization, similar to that observed in microcephaly **(Figure 7I and S11).**

## Discussion

Using a cell-type-specific gene enrichment analysis of human brain organoids, we identified *KIAA0408* (CCB79), a previously uncharacterized vertebrate-specific gene, and assigned its function as a primate-specific factor involved in cilium initiation. Our approach demonstrates the effectiveness of using human organoids to identify primate- and human-specific gene functions that contribute to the expansion of the human neocortex^60,61^. Molecularly, CCB79 is a centriolar DAP and interacts with other DAP components, potentially forming a biochemical cilia assembly complex critical for cilium initiation **(Figure 2)**. DAPs anchor the mother centriole to the plasma membrane during ciliogenesis. In this process, CCB79 is a cell cycle-regulated molecule. It appears in DAP only at the onset of ciliogenesis (G1/G0 phase), where it recruits TTBK2, another DAP component, to initiate ciliogenesis **(Figures 2, 4 and S3).**

Although centriolar DAPs and cilia are typically expressed across all neural cell types, CCB79’s expression is notably higher in intermediate progenitors, which are relatively abundant neurogenic NPCs in gyrencephalic brain species critical for generating neurons in the evolutionary expansion of the human cortex^9,10^. This suggests that context-specific ciliary functions may have evolved during brain development, warranting further research into the roles of cilia in human brain development and disease.

Notably, although CCB79 is conserved in vertebrates, the human gene has experienced significant changes and faced an evolutionary challenge in initiating ciliogenesis **(Figure 3).** Our rescue experiments show that the mouse gene neither targets the ciliary base nor restores ciliogenesis in human iPSCs lacking CCB79 **(Figure 3C-D)**. This suggests that human CCB79 has specifically evolved to promote ciliogenesis in human cells, leading us to hypothesize that human cilia may have recently evolved, at least in specific cell types of the developing brain, in terms of composition, length, and dynamics. This is surprising because centrioles containing DAPs are very ancient structures conserved from single-celled organisms to humans; thus, significant evolutionary changes in them are unexpected. However, it appears that CCB79 has undergone rapid evolution in terrestrial animals. It will be interesting to investigate the rodent ortholog’s function and test the expression of human CCB79 in rodents and determine whether it can alter cilia, thereby affecting cell composition and inducing gyrification in lissencephalic rodent brains.

Our findings indicate that telencephalic and ventral progenitors, as well as the inhibitory interneurons they generate, are particularly susceptible to CCB79 disruptions **(Figure 5)**. We could precisely identify these defects because our organoids consistently mimic the in vivo cell type diversity of human brain development, including essential progenitor populations and both excitatory and inhibitory interneurons derived from the medial ganglionic eminence.

To date, no non-syndromic ciliopathy or neurodevelopmental disorder has been linked to CCB79. However, our findings suggest that the CCB79 gene might be a risk factor for microcephaly and autism-like disorders. This is because most affected cell types after CCB79 perturbation are ventral progenitors, which typically develop into GABAergic interneurons and oligodendrocyte precursors under normal conditions. When CCB79 is perturbed, the fate determination of ventral progenitors is disrupted, leading to the formation of oligodendrocyte precursors in an unscheduled manner at the expense of ventral progenitors. Earlier studies have shown that prolonged or excessive SHH causes ventral progenitors to differentiate into oligodendrocytes, although the underlying mechanism remains unclear. Our data address this gap, as CCB79-KO cells lacking functional cilia display signs of excessive SHH activation, which may be a key factor in disrupting the balance.

Our study has limitations. First, we were unable to generate homogeneous populations of mouse transcript-expressing iPSCs with regulated expression of the gene product. This is primarily because expression of CCB79 is temporarily regulated and appears only at the onset of cilium initiation **(Figure S2D).** Future studies should investigate the expression of the human transcript in the mouse endogenous locus or the mouse gene knocked into human iPSCs to dissect the specific aspects of cilia and progenitor biology affected by the mouse and human genes. Secondly, we do not know if CCB79 mutations have occurred in human patients. It would be interesting to screen patient cohorts for CCB79 mutations and determine if CCB79 is a risk factor for neurodevelopmental disorders.

Nevertheless, our findings provide new insights into cilia biology and its previously unknown connections to the maintenance of progenitors and human brain development mechanisms **(Figures 5-7 and S11).** Human cortical development involves intricate processes that generate a wide variety of cell types, with their diversity and connections carefully regulated through a complex interplay of transcription, epigenetics, cell adhesion, and polarity factors. CCB79 removal, however, primarily affects DAP assembly and ciliogenesis, yet it significantly disrupts cell diversity, indicating a crucial role for cilia in regulating brain organization. In terms of cilia biology, since CCB79 has never been studied and is a novel DAP component, our discoveries will improve the molecular understanding of DAP roles in cilium initiation. The results and resources presented lay the groundwork and open new avenues for cilia research to investigate CCB79’s potential isoforms, interaction partners, and regulatory mechanisms in ciliogenesis. Additionally, these findings strengthen the underexplored connection between cilia dynamics and human brain development.

## Acknowledgment

This work is supported by Fritz Thyssen Foundation (10.20.2.031MN), and Deutsche Forschungsgemeinschaft (DFG, German Research Foundation) – Project-ID 503306912 – FOR5547, DFG, GO 2301/5-2.

## Author contributions statement

JG conceived the concept, and DR, VJ, and SW equally contributed to the work, with DR performing most of the imaging experiments. VJ performed the transcriptomic data analysis, and SW conducted the initial localization findings and was involved in gene discovery. EG reprogrammed iPSCs and generated NPC experiments; OSV conducted 3D tissue clearing and light sheet imaging; SA and Ml reprogrammed cells; ZZ, AM, TB, and CG conducted SIM and STED-microscopy imaging; EC, EC, NP and TB conducted proximal proteomics, MS analysis and Western blot experiments; NA edited the iPSCs and ablated CCB79; MR and GC conducted EM experiments. JG, EG, DR, and VJ wrote the manuscript. All authors have read the manuscript and approved it for publication.

## METHOD DETAILS

### Human iPS cells and culturing

Human iPS cell lines IMR90 and BJ151 were cultured in mTeSR^™^ plus medium (Stem Cell Technologies) on Matrigel-coated dishes (Corning) at 37°C with 5% CO_2._ They were routinely checked for mycoplasma contamination using the MycoAlert Kit (Lonza). Cells were dissociated into small aggregates with ReLeSR (Stem Cell Technologies) every 5-7 days and split 1:5 onto new Matrigel-coated dishes. Unless otherwise noted, most analyses were performed using organoids and NPCs derived from IMR90 (Coriell) or BJ151, which was reprogrammed in-house. BJ151 fibroblasts were expanded and electroporated using the Lanza Nucleofector 4D with Yamanaka factors delivered via EBNA1/oriP episomal vectors expressing six reprogramming factors (Oct4, Sox2, Klf4, L-Myc, Lin28, and p53 shRNA). After electroporation, fibroblasts were maintained in fibroblast medium for 5 days, then transitioned to N2B27/FGF medium for 10 days, followed by 5 days in Essential-8 medium. By day 20, three clones were selected and plated on pre-coated wells with rhLaminin-521 in Essential-8 medium. The generated iPSCs were further passaged multiple times and tested for pluripotency via FACS analysis. Successful reprogramming was confirmed based on colony morphology, characterized by defined borders and a high nucleus-to-cytoplasm ratio, as well as dual positivity for the human pluripotency markers TRA-1-60 and TRA-1-81.

### Generation of CCB79 (KIAA0408) knock-out iPSCs

Two single guide RNAs (sgRNAs) targeting exon 3 of the *KIAA0408* gene were chosen based on high predicted on-target activity and minimal off-target effects: sgRNA1: ATGTCGGTCGATGTGCAGCA, and sgRNA2: CATCGACCGACATCAGATCC. Each sgRNA was cloned into a lab-modified pX330A-derived plasmid encoding *S. pyogenes* Cas9, a U6 promoter-driven sgRNA cassette, and a GFP reporter to monitor transfection efficiency. Oligonucleotides were synthesized with BbsI-compatible overhangs as follows: sgRNA1-FP: CACCGATGTCGGTCGATGTGCAGCA, sgRNA1-RP: AAACTGCTGCACATCGACCGACATC; sgRNA2-FP: CACCGCATCGACCGACATCAGATCC, and sgRNA2-RP: AAACGGATCTGATGTCGGTCGATGC.

Golden Gate cloning was used to insert phosphorylated, annealed oligonucleotides into the BbsI-digested vector backbone. Briefly, oligonucleotides were phosphorylated using T4 Polynucleotide Kinase (NEB) at 37°C for 30 minutes and then annealed by gradual cooling from 95°C to 25 °C. The pX330A plasmid was linearized by digestion with FastDigest BbsI (Thermo Fisher) and purified by column purification. Annealed oligos (1:200 dilution) were ligated into the digested vector using T4 DNA Ligase (NEB) in a Golden Gate assembly. Ligation products were treated with PlasmidSafe DNase (EpiCentre) to remove linear or un-ligated DNA and subsequently transformed. Positive clones were confirmed by Sanger sequencing, and validated constructs were used for downstream transfection.

hiPSC IMR90 cells were cultured on Matrigel-coated plates in mTeSR1 medium and dissociated using Accutase. For each sgRNA, 75,000 cells were seeded per well in a 12-well plate. Cells were transfected using GeneJuice (Sigma-Aldrich) with 1 µg of plasmid DNA per well. After 24 hours, the cells were treated with 1 μg/mL puromycin for an additional 24 hours. Transfection efficiency was assessed by monitoring the fluorescence of GFP.

Targeted mutations revealed by the mismatch-sensitive T7 endonuclease I (T7E1, NEB) assay. For clonal validation, single cells were seeded at low density to allow colony formation. Individual clones were manually picked, expanded, and screened by PCR and Sanger sequencing to confirm the presence of indels or frameshift mutations introduced by CRISPR-Cas9 editing.

### NPC generation

NPCs were differentiated from iPSCs as previously described ^38–40^. iPSCs were dissociated into single cells using Accutase. For the neurospheres, 45,000 iPSCs were seeded into a 96-well V-bottom plate and maintained in NIM for 5 days. Neurospheres were transferred into a PLO/Laminin-coated 35 mm dish using a 200 μL cut tip and evenly distributed. The dish was incubated at 37 °C with 5% CO₂ for 7 days, with a medium change performed daily. The adherent neurospheres with neural rosettes were washed once with DMEM-F12 and incubated for 1 hour in Neural Rosette Selection Medium at 37 °C and 5% CO₂. After incubation, DMEM-F12 was used to dislodge only the neurospheres from the dish. The neurospheres were gently resuspended using an uncut 1000 μL pipette tip to generate smaller aggregates. These aggregates were transferred to a Falcon tube, centrifuged at 700 × g for 3 minutes, and resuspended in 1 mL of NIM supplemented with 10 μM ROCK inhibitor. The cell suspension and small aggregates were evenly distributed in a PLO/Laminin-coated 35-mm dish and incubated at 37°C with 5% CO₂. For expanding the NPCs, medium changes were performed with STEMdiff™ Neural Progenitor Medium (STEMCELL Technologies, Cat #05833) on a daily or every-other-day basis until confluency was achieved.

### Brain organoid generation

The brain organoids were generated by a non-directed differentiation method as previously described ^35^. The iPSCs were cultured in mTeSR Plus culture medium and passaged using ReLeSR. The cells were regularly checked for mycoplasma contamination and their stem cell morphology. For initiating neural differentiation, cells were dissociated into single cells. Therefore, iPS cells were carefully washed once with pre-warmed DMEM/F12 (GIBCO) and then treated for 5 minutes at 37°C with Accutase (Sigma-Aldrich). After cell collection, centrifugation, and counting, cells were diluted in neural induction medium (NIM, Stem Cell Technologies) plus Rock inhibitor (10 mM, Y-27632 2HCl, Selleck Chem) at a concentration of 3,5 × 10^5 cells/mL. 100 μL of this cell suspension was added to each well of a 96-well V-bottom low-adherent plate and incubated at 37°C and 5% CO2. Half of the medium was changed every day for 5 days until neurospheres were formed.

### Neurospheres in Petri dishes and spinner flasks

At day 5 of neural induction, the neurospheres were carefully collected using a 200 μL pipette tip and embedded into Matrigel basement membrane extracted from the Engelbreth-Holm-Swarm (EHS) mouse sarcoma. The embedded neurospheres were cultured in a petri dish for another 4 days, containing neurosphere medium. On day 9, the neurospheres were transferred into a spinner flask containing brain organoid medium and were then cultured for an additional 14 days. Starting from the first day of differentiation, the process takes a total of 24 days.

### Immunofluorescence microscopy of 2D cells and 3D brain organoid tissues

Human iPSCs, NPCs, and brain organoids were immunostained using the following method. Materials were fixed in 4% PFA (Merck Millipore) for 30 min at room temperature (RT) and treated with PBS-Glycine (0.225 g Glycine per 100 mL PBS). For organoids, PBS-Glycine was removed, and 30% Sucrose (Merck Millipore) was added to dehydrate the tissues overnight at 4°C. The next day, organoids were embedded into Tissue-Tek® Cryomold® Cryomolds (Science Services) containing Tissue-Tek® O.C.T. Compound solution (Science Services) and immediately stored at −80°C for a minimum of 24 hours. Frozen specimens were embedded and cryo-sectioned at 12 μm thickness with Cryostat (Leica CM3050 S) and placed onto Poly-L-lysine (Sigma-Aldrich) pre-coated Superfrost glass slides (0.01% PLL diluted in distilled H2O and coated for 5 min at RT, subsequently dried for 3 hours at RT) (Thermo Scientific). Slides with organoid sections were dried for 1 hour at RT and then kept for long-term storage at −80°C.

Before immunofluorescent staining procedures, we thawed the slides with organoid sections for 30 min at room temperature, washed 2x 3 min with PBS-Glycine (0.225 g per 100 ml) and permeabilized for 15 min in 0.5% Triton X-100/0.1% Tween (Sigma Aldrich) in PBS solution at RT, and washed again 2x 3 min with PBS-Glycine. Subsequently, sections were blocked with blocking solution (0.5% Fish gelatin in PBS) for 1 hour at RT. Primary antibodies were diluted in blocking solution, and sections were incubated with primary antibodies for 2 hours at RT.

2D cells were treated on glass coverslips fixed with either 4% PFA or ice-cold methanol. Methanol fixation was used to stain CCB79. After three washes with blocking solution, the corresponding secondary antibodies (goat-anti-mouse Alexa 488 or 647; donkey-anti-rabbit Alexa 488, 594, or 647; donkey-anti-sheep 647; or donkey-anti-goat Alexa 594) were diluted in blocking solution (1:800) and incubated for one hour at room temperature protected from light. Finally, sections were stained with DAPI, washed twice for three minutes with blocking solution, and after a final wash with distilled water, briefly air-dried, and mounted with Mowiol (Sigma Aldrich) and glass coverslips. Slides were dried for one hour at room temperature and then stored at 4°C in the dark until imaging. For miniTurbo experiments, cells were fixed with 100% methanol at −20 °C. To verify the biotinylation, all samples were stained with a 1:200 dilution of streptavidin-488 and anti-cep152 overnight at 4 °C. The secondary antibodies were then incubated for one hour at room temperature.

### Confocal and light-sheet microscopy

Tissue sections or 2D cells on glass coverslips were imaged using a Leica SP8 confocal microscope. Our experiments utilized laser lines at 405, 488, 561, and 633 nm and were imaged using an objective lens with a 63x oil-immersion magnification. For light-sheet imaging, the brain organoids were fixed in 4% warm PFA for 1 hour at room temperature on a gentle rocker. The samples were then washed twice in PBS before being stained with antibodies. Briefly, the organoids were permeabilized in a permeabilization buffer for 1 hour at room temperature, followed by blocking for 1.5 hours in a blocking solution with fish gelatin. The organoids were incubated overnight at 4°C in the blocking solution with primary antibodies against SOX2 (Abcam, 1:400 dilution) and MAP2 (Proteintech, 1:400 dilution). The organoids were then washed three times in blocking solution for 30 minutes. Next, the organoids were incubated with a primary antibody against ARL13B (BiCell Scientific, 1:400 dilution) for 3 hours at room temperature. Afterward, the organoids were washed as previously described, then incubated with secondary antibodies (1:1000 dilution) overnight at room temperature. The following day, the organoids were washed again as described and embedded in low-melting agar.

The embedded organoids were sequentially dehydrated using 30%, 50%, 70%, 90% and absolute ethanol. After the dehydration step, the blocks were treated with Ethyl Cinnamate overnight, followed by imaging.

Imaging data acquisition was performed using the Light Sheet 7 system (Carl Zeiss) with a 5x objective at a 1x zoom and a 20x objective at a 2x zoom. The same step size (z step thickness) and imaging depth (number of z steps) were used for generating maximum intensity projections, sections of the VZs, volumetric rendering, and image-based quantifications. Raw data underwent channel alignment (case-specific) in Zen Black software (Zeiss) before analysis. Data analysis was conducted with Arivis Pro (Zeiss). Image-based machine learning and segmentation techniques were employed to quantify the volume distribution of VZ, lumen diameters, and the number of rosettes across samples. The blob finder module facilitated segmentation and quantification of progenitor densities within fixed volumes in the rosettes across samples. Data were exported to MS Excel. Statistical analyses and graph plotting were performed using GraphPad Prism 10. The raw images were processed with ImageJ, Adobe Photoshop 2024, and Adobe Illustrator 2024. Movies of organoid Z-stack images were processed using ZEISS arivis Pro.

### STED (Stimulated Emission Depletion) Super-Resolution Imaging

Two-color STED images were obtained using the Abberior Expert Line, equipped with pulsed excitation and depletion lasers, along with an Olympus UPlanSApo 100x/1.4 oil immersion objective as previously described^62^. Proteins of interest were labeled with Abberior Star-Orange (Distal Appendage Protein FBF1 and Cilia Marker ARL13B). Abberior Star-Red (rat CCB79) dyes, attached via secondary antibodies, were excited by 561 nm and 640 nm pulsed diode lasers, respectively. The fluorescence signals were suppressed by a pulsed 775 nm depletion laser (total power = 3 W). DAPI was excited using 405 nm and 488 nm pulsed diode lasers. Before acquiring images, the excitation and depletion beams were aligned for both 2D and 3D STED. Briefly, the centers of the excitation and depletion beams were initially overlapped by scanning 150 nm gold beads (BBI Solutions) in reflection mode. Then, TetraSpeck beads of four colors (TetraSpeckTM Microspheres, 100 nm, fluorescent blue/green/orange/dark red) were used to correct mismatches between scattering and fluorescence modes. To ensure accurate positioning of the same beads imaged by different laser lines, individual confocal and STED channels were compared and adjusted on the Imspector SLM (spatial light modulator) adjustment panel as the final step. During STED imaging, sequential line-by-line scanning minimized photobleaching, starting with the Star-Red channel, followed by the Star-Orange channel. Images were acquired according to the Nyquist criterion for super-resolution, with a pixel size of 20 nm and a dwell time of 10 µs. A 100 nm distance was maintained between slices in the XYZT stacks. The raw STED images were deconvolved using Huygens Professional version 24.10 (Scientific Volume Imaging, https://svi.nl/) with the classic maximum likelihood estimation (MLE) algorithm, incorporating lateral drift stabilization, over 27 iterations.

### Structured Illumination Microscopy (SIM) Super-Resolution Imaging

To provide complementary super-resolution imaging with lower phototoxicity and a broader field of view, SIM was performed to image the same protein-labeled samples used for STED. Images were acquired using a commercial Zeiss Elyra 7 Lattice SIM system, equipped with a Plan-Apochromat 100x/1.46 oil immersion objective, and sensitive sCMOS cameras. Structured illumination was applied in a 3D mode, and image reconstruction was performed using Zeiss ZEN Black, with default parameters for modulation contrast and noise filtering. SIM images were acquired at Nyquist sampling conditions, with voxel sizes ranging from 40 to 60 nm in the xy plane and from 120 to 150 nm in the z direction. Image post-processing was performed using Fiji.

### Antibodies used

The following antibodies were used. Rat anti-CCB79 (1:100, generated in house via BiCell Scientific), Rabbit anti-CCB79 (1:500, generated in house via BiCell Scientific), Rabbit anti-CEP164 (1:200, Proteintech), Rabbit anti-TTBK2 (1:100, Sigma Aldrich), Rabbit anti-SCLT1 (1:25, Proteintech), Rabbit anti-FBF1 (1:200, Proteintech), Rabbit anti-CEP89 (1:500, Proteintech), Rabbit anti-CEP83 (1:300, Sigma Aldrich), Rabbit anti-ARL13B (1:100, Proteintech), Rat anti-ARL13B (1:100, BiCell Scientific), Mouse anti-ARL13B (1:200, Proteintech), Mouse anti-GT335 (1:2000, AdipoGen), Rab anti-CP110 (1:200, Proteintech), Mouse anti-CPAP (1:25, in house), Rabbit anti-ODF2 (1:100, Proteintech), Mouse anti-SOX2 (1:500, Abcam), Rabbit anti-PAX6 (1:10, Proteintech), Rab anti-FoxG1 (1:300, Abcam), Mouse anti-Nestin (1:100, Novus Biologicals), Rabbit anti-TUJ1 (1:300, Sigma-Aldrich), Rabbit anti-pVIM (1:500, Abcam), Rabbit anti-TBR2 (1:100, Abcam), Rabbit anti-PTPRZ1 (1:200, Sigma-Aldrich), Rabbit anti-MAP2 (1:200, Proteintech). We used Alexa Fluor Dyes conjugated with goat/donkey anti-mouse, anti-rabbit, or anti-rat antibodies (1:1000, Molecular Probes, Thermo Fisher, USA) as secondary antibodies. In addition, DAPI at 1 μg/ml (Sigma Aldrich, USA) was used to stain the nucleus.

### Tissue-specific expression analysis of CCB79 (KIAA0408)

was conducted using data from the UCSC Genome Browser GTEx Gene V8 Track (Human genome GRCh38/hg38), which summarizes mean expression across fetal and adult human tissues. The data were loaded into pandas and organized into separate fetal and adult expression matrices. To facilitate comparison, expression values were normalized using min–max scaling based on the fetal expression range as the reference. Heatmaps created with seaborn display tissues sorted by fetal KIAA0408 expression, using a consistent viridis color scale for both fetal and adult panels to emphasize developmental differences. Single-cell RNA-seq data from the human developing neocortex were sourced from a publicly available dataset^38^that includes cells from 20 distinct developmental stages. Expression levels of genes, including SOX2, PAX6, DCX, and CCB79 (KIAA0408), were visualized across these stages using a dotplot in Scanpy, with standardized expression to illustrate gene dynamics during brain development.

### Organoid dissociation, single-cell preparation, and scRNA sequencing

Human iPSC-derived brain organoids at Day 14 from both control and CCB79-KO genotypes were processed for single-cell preparation using a modified protocol adapted from ^55^. Briefly, 3–4 intact organoids were rinsed three times with 1× PBS to remove residual media. Organoids were then incubated in 2 mL Accutase (StemPro) supplemented with 0.2 U/µL DNase I (Roche, [Catalog #]) for approximately 45 minutes at 37°C, with gentle flicking every 10 minutes to facilitate dissociation into single cells. If visible aggregates or debris remained after dissociation, the suspension was sequentially passed through 40 µm strainers to ensure a uniform single-cell suspension. The dissociated cells were pelleted by centrifugation at 900 × g for 3 minutes, washed once with 1 mL of 1× PBS, and then recentrifuged under the same conditions. Cell viability and concentration were assessed using Trypan Blue exclusion on a Countess™ Automated Cell Counter (Thermo Fisher Scientific).

Single-cell fixation was performed according to the 10x Genomics Flex Gene Expression Profiling protocol. Briefly, up to 2 million cells were pelleted at 350 × g for 5 minutes at 4°C, then resuspended in 1 mL of Fixation Buffer and incubated for 16–24 hours at 4°C. Fixation was quenched by centrifugation at 850 × g for 5 minutes at room temperature, followed by the addition of 1 mL Quenching Buffer (1× Conc. Quench Buffer; 10x Genomics PN-2000516). To enable long-term storage, 0.1 volumes of Enhancer (10x Genomics PN-2000482) and 10% glycerol were added, and samples were stored at −80°C. Before library preparation, samples were thawed at room temperature, centrifuged at 850 × g for 5 minutes, and resuspended in 1 mL of 0.5× PBS containing 0.02% BSA.

Hybridization of probe sets was carried out according to 10x Genomics recommendations. Up to 2 million fixed cells were used per hybridization reaction in a total volume of 80 µL, consisting of 60 µL hybridization buffer and 20 µL of either Human Transcriptome Probe Panel v1. Hybridizations were incubated at 42°C for 16–24 hours. After hybridization, samples were diluted in Post-Hybridization Wash Buffer and analyzed using an automated Cell Counter.

For standard pooling, equal cell numbers from each hybridization were combined; for differential pooling strategies, cell numbers were adjusted as needed. Cell pools underwent three washes in Post-Hybridization Wash Buffer at 42°C for 10 minutes each, followed by resuspension in Post-Hybridization Resuspension Buffer. Samples were filtered through a 30 µm filter to remove aggregates, then recounted before loading on the Chromium system.

NGS library preparation was performed using the 10x Genomics multiplexed Fixed RNA Profiling protocol, following the manufacturer’s guidelines (CG000527 and CG000478 for sample fixation). Libraries were prepared with input ranging from approximately 140,000 to 2,000,000 cells per sample. Sixteen samples were pooled into a single batch, aiming for an average of 8,000 cells per sample, and amplified through 20 PCR cycles. A post-PCR purification step was conducted to remove residual primers and adapter dimers. Library quality was assessed using a High Sensitivity DNA chip on a 2100 Bioanalyzer (Agilent Technologies) and quantified with the Qubit 1X dsDNA HS Assay Kit on a Qubit 4.0 Fluorometer (Invitrogen by Thermo Fisher Scientific). Sequencing was carried out on a NextSeq 2000 platform with a P3 flow cell, using 28 cycles for read 1, 90 cycles for read 2, and 10 cycles each for i7 and i5 indexes at the Core Facility Genomics, Institute of Molecular Biology GmbH (IMB), Mainz, Germany.

### Data processing and analysis

#### Cell Ranger Analysis

Raw 10x Genomics Flex data (FASTQ files) were processed using Cell Ranger v7.1.0 (10x Genomics) and aligned to the human reference (GRCh38-2020-A, provided by 10x Genomics) using the Cell Ranger multi-pipeline with default parameters. The pipeline performed demultiplexing, read alignment, barcode processing, and UMI counting to generate feature-barcode matrices for downstream single-cell RNA-seq analysis. For downstream single-cell RNA-seq data processing, highly variable genes were identified, and the total number of counts per cell, along with the percentage of mitochondrial reads, were regressed out to normalize technical variation. Principal component analysis (PCA) was performed, followed by batch correction using the Harmony package (v1.2). To account for local neighborhood structure, a k-nearest neighbor graph was computed using the BBKNN algorithm, and a two-dimensional representation was generated with the UMAP and Force Atlas2 (FA) algorithm for visualization. Cell type annotation for the CCB79-KO and control was performed using canonical marker genes and top differentially expressed genes (DEGs) identified for each cluster **(Supplementary Table 3)**. The proportion of each annotated cell type was used as input for principal coordinate analysis (PCA) to assess inter-sample relationships. Unsupervised clustering was performed using the Leiden algorithm, and marker genes for each cluster were identified through differential expression analysis using the Wilcoxon Rank Sum test and t-tests. Pseudotime trajectory inference was conducted using the diffusion pseudotime (DPT) algorithm, with proliferative radial glia cells designated as the root state. All analyses were implemented in Scanpy (version ≥ 1.10), and the whole pipeline was scripted and logged to ensure reproducibility.

### Bulk RNA transcriptomic analysis

Bulk RNA sequencing analysis was conducted on six samples, representing two experimental groups (CCB79-KO and control), with three biological replicates each. Metadata detailing sample identifiers, condition labels, and replicate numbers was compiled into a structured tab-delimited file for downstream processing and analysis. Briefly, raw FASTQ files were aligned to the human reference genome (GRCh38) using the STAR aligner (Dobin et al., 2013) version 2.7. Gene-level quantification was then performed using feature counts from the Subread package version 2.0.2, using the corresponding annotation file. The resulting count files were parsed to extract gene-level expression values and merged into a unified expression matrix across all samples ^63,64^. The matrix was subset to include only protein-coding genes, and genes with fewer than 10 total counts across all samples were excluded to reduce technical noise.

Differential expression analysis was conducted using the DESeq2 package (^65^version 1.44.0. All analyses compared CCB79-KO to control. Significantly differentially expressed genes (DEGs) were identified based on an adjusted p-value threshold of 0.05, corrected with the Benjamini-Hochberg method. For visualization, MA plots and volcano plots were created using DESeq2 and ggplot2 version 3.5.2, while heatmaps of the top 50 DEGs were generated with the pheatmap package version 1.0.13. Functional enrichment analyses used the clusterProfiler package ^66^, version 4.12.6, with a focus on Gene Ontology (GO) biological processes and KEGG pathways. Significant DEGs were mapped from gene symbols to Entrez Gene IDs using the org. Hs.eg.db annotation database version 3.19.1, and enrichment results were visualized as dot plots. All analyses were carried out in R (version 4.0 or later), with the complete pipeline scripted and logged for reproducibility.

### Hedgehog signaling activity

To assess Hedgehog (HH) signaling activity, we implemented a weighted pathway activation scoring strategy using z-transformed expression values of genes associated with key functional roles, similar to previously described contexts that use gene expression of signaling components to infer pathway activation^67–70^. Briefly, genes were categorized into three main functional groups: activators, suppressors, and ligands. Activators (e.g., GLI1, GLI2, SMO, BCL2, WNT2B) were assigned a weight of +1, reflecting their positive regulatory role in HH signal transduction and transcriptional output ^71,72^. Suppressors (e.g., PTCH1, PTCH2, SUFU, HHIP) were given a weight of −1, as they inhibit pathway progression by sequestering or repressing activators ^73^. Ligands (e.g., SHH, IHH, DHH) were assigned a moderate weight of +0.5 to reflect their role as extracellular initiators of the pathway without directly encoding for downstream transcriptional regulators. The weighted HH score was computed by first scaling gene expression across samples (z-transformation), then multiplying each gene’s expression value by its assigned weight and finally summing these weighted scores for each sample. This composite metric estimates the net activation state of the HH pathway in each biological replicate, capturing the balance between activation and repression signals at the transcriptomic level.

To examine the biological significance of HH pathway modulation, we then explored its relationship with the expression of genes involved in forebrain development. The HH signaling pathway plays a vital role in early brain patterning, including the regionalization of the forebrain, midbrain, and hindbrain, with a particular emphasis on ventral forebrain specification^74–76^. Genes such as FOXG1, PAX6, TBR1, DLX2, and EMX2 are key regulators of telencephalon development and neuronal differentiation. Additionally, DLX2 and GSX2 are recognized as transcriptional targets or downstream effectors of the HH pathway in neural progenitors^77^. To explore this connection, we correlated the weighted HH scores with the expression levels of selected forebrain marker genes across control and CCB79-KO samples using Pearson correlation. These correlations were visualized through heatmaps and bar plots to identify candidate genes that may be co-regulated or influenced by HH pathway activity. The findings offer insights into whether changes in HH signaling in the CCB79-KO condition could affect forebrain-related transcriptional programs, potentially reflecting neurodevelopmental dysregulation linked to genetic perturbations.

### Transmission electron microscopy (TEM)

iPSCs were washed once in PBS, then fixed overnight at 4°C in 2.5% glutaraldehyde and 1.8% sucrose in PBS. Samples were rinsed in PBS and post-fixed with 1% osmium tetroxide for 1 hour at 4°C. After fixation, the samples were dehydrated in a graded ethanol series and embedded in Epon-Araldite resin for 48 hours at 60°C. Ultrathin sections (40-60 nm) were prepared using the Reichert Ultracut E ultramicrotome. These sections were stained with uranyl acetate and lead citrate and observed with a FEI Tecnai G2 Spirit 255 transmission electron microscope operating at 100 kV.

### Molecular biology, cloning, and lentiviral generation

The cDNA sequences of the human or mouse CCB79 gene were cloned into the PiggyBac plasmid pXLONE-MCS-GFP. Primers incorporating KpnI and NotI restriction sites were designed to amplify the target cDNA. Both the PCR product and the plasmid were digested with KpnI and NotI, followed by ligation. The ligation mixture was transformed into TOP10 competent *E. coli* and plated on LB agar containing ampicillin. Positive clones were screened by restriction digestion and confirmed by Sanger sequencing to verify the correct insert was present. MiniTurbo and CCB79-miniTurbo were cloned into the pCW57-GFP-2A-MCS plasmid (Addgene: #71783).

HEK293FT cells were used as a host for generating lentiviral particles. Cells were co-transfected with psPAX2 (addgene #12260), pMD2.G (addgene #12259), and pCW57-MiniTurbo or pCW57-CCB79-miniTurbo plasmids. Cell medium collected from transfected HEK93FT cells and concentrated with PEG over three days. Concentrated virus transduced to HTET-RPE1 cells. Transduced cells were selected with 10 μg/mL puromycin.

### Western blot

Cell pellets were lysed in a standard lysis buffer on ice. 60-70 µg of protein samples were resolved on 10% or 12% acrylamide gels and transferred to nitrocellulose membranes. The membranes were then blocked with 5% skim milk at room temperature for 1 hour. All primary antibodies were incubated overnight at 4 °C, and secondary antibodies were incubated at room temperature for 1 hour. SuperSignal Western Pico or Femto was used to detect the signal. Antibody dilutions were as follows: anti-rabbit CCB79 (C-terminal), anti-mouse beta-actin (1:3000), and peroxidase-conjugated secondary antibodies at 1:3000 (Life Technologies).

### Biotinylation and miniTurbo-based affinity purification and preparation for liquid chromatography tandem mass spectrometry (LC-MS/MS)

#### Pull-down of biotinylated proteins

RPE-miniTurbo and RPE-CCB79-miniTurbo cells were incubated with 3 μg/mL doxycycline for 72 hours. (doxycycline renewed every day to ensure expression). After induction, cells were treated with 500 μM d-biotin (B1595) for 30 minutes. After 30 minutes, the cells were washed with PBS, and samples were prepared according to Cirri et al. 2025 ^78^ with some modifications. Briefly, four biological replicates, 8 million cells each (corresponding to approximately 0.8 mg protein) were resuspended in 500 µl lysis buffer (50 mM Tris pH 7.5; 150 mM NaCl; 1 mM EDTA; 1mM EGTA; 1% (v/v) Triton 0; 1 mg/ml aprotinin (Carl Roth); 0.5 mg/ml leupeptin (Carl Roth); 250 U turbonuclease (MoBiTec GmbH); 0.1% (w/v) SDS), respectively, and incubated for one hour at 4°C, then sonicated for 10x 60sec on/30sec off using a Bioruptor (Diagenode, Beligum) and then spun down (17000g) for 20 minutes at 4°C.

For the acetylation of the streptavidin cartridges and the subsequent loading of the lysates onto the liquid handler Bravo AssayMAP, the protocol was the same as in Cirri et al. (2025 ￼)^78^, modified as follows: Cartridges were equilibrated with 200 µl PBS (at a rate of 10 µl/min). For each replicate, 50 µl 10 mM sulfo-NHS acetate was used. The reaction was set to a 6 µl volume and incubated at 25°C for 30 min. The reaction chase was 100 µL with a flow rate of 10 µL/min. Before loading, cartridges were equilibrated using Internal Cartridge Wash 1, with 200 µl of lysis buffer (20 µl/min). Before and after equilibration, Cup Wash was used with default settings. Samples were loaded at a speed of 10 µl/min. No on-bead digestion was performed; instead, biotinylated proteins were eluted twice with 15 µL of 10% TFA in acetonitrile (90 µL/min). Peptides were eluted in two steps (1 x 25 µl and 1 x 50 µl) using 50 mM AmBic (no internal cup wash, 10 µl/min). Trypsin (0.5 µg, Promega) was added to the elution and incubated overnight at 37 °C. Biotinylated peptides were eluted twice with 15 µl 10% TFA in acetonitrile (90 µl/min). The syringe was washed with 150 µl 20% acetonitrile. Biotinylated peptide elution was dried and reconstituted in 50 µl 50 mM HEPES. The pH was tested and adjusted with sodium hydroxide to pH 6 −8.0. The eluate was digested with 0.5 µg of trypsin at 37°C overnight. Both elutions were cleaned up using Waters Oasis® HLB μElution Plate 30 μm (Waters) according to the manufacturer’s instructions. The eluates were dried using a speed vacuum centrifuge (Eppendorf Concentrator Plus, Eppendorf AG, Germany) and stored at −20° C. Before analysis, samples were reconstituted in in MS Buffer (5% acetonitrile, 95% Milli-Q water, with 0.1% formic acid), spiked with iRT peptides (Biognosys, Switzerland) and loaded on Evotips (Evosep) according to the manufacturer’s instructions. In short, Evotips were first washed with Evosep buffer B (acetonitrile, 0.1% formic acid), conditioned with 100% isopropanol and equilibrated with Evosep buffer A. Afterwards, the samples were loaded on the Evotips and washed with Evosep buffer A. The loaded Evotips were topped up with buffer A and stored until the measurement.

#### Data Acquisition and Processing for DIA Samples

Peptides were separated using the Evosep One system (Evosep, Odense, Denmark), equipped with a 15 cm × 150 μm i.d. column packed with a 1.5 μm Reprosil-Pur C18 bead (Evosep Performance, EV-1137, Denmark), heated at 45 °C. The samples were run with a pre-programmed proprietary Evosep gradient of 44 minutes (30 samples per day), using water and 0.1% formic acid as solvent A, and solvent B, consisting of acetonitrile and 0.1% formic acid. The LC was coupled to an Orbitrap Exploris 480 (Thermo Fisher Scientific, Bremen, Germany) using PepSep Sprayers and a Proxeon nanospray source. The peptides were introduced into the mass spectrometer via a PepSep Emitter with an outer diameter of 360 μm and an inner diameter of 20 μm, heated to 300°C, and a spray voltage of 2 kV was applied. The injection capillary temperature was set at 300°C. The radio frequency ion funnel was set to 30%. For DIA data acquisition, full-scan mass spectrometry (MS) spectra with a mass range of 350–1650 m/z were acquired in profile mode using the Orbitrap at a resolution of 120,000 FWHM. The default charge state was set to 2+, and the filling time was set to a maximum of 20 ms, with a limitation of 3 × 10^^6^ ions. DIA scans were acquired with 40 mass window segments of differing widths across the MS1 mass range. Higher collisional dissociation fragmentation (normalized collision energy 30%) was applied, and MS/MS spectra were acquired with a resolution of 30,000 FWHM with a fixed first mass of 200 m/z after accumulation of 1 × 10^6^ ions or after filling time of 45 ms (whichever occurred first). Data were acquired in profile mode. For data acquisition and processing of the raw data, Xcalibur 4.0 (Thermo) and Tune version 4.0 were used.

#### Data Analysis for DIA Samples

DIA raw data were analyzed using the directDIA pipeline in Spectronaut v.19 (Biognosys, Switzerland). The data were searched with the following modifications: Oxidation (M), Acetylation (Protein N-term), and biotinylation (Biotin_K). A maximum of 2 missed cleavages for trypsin and five variable modifications were allowed. The identifications were filtered to achieve an FDR of 1% at both the peptide and protein levels. Relative quantification was performed in Spectronaut using the LFQ QUANT 2.0 method with Global Normalization, precursor filtering percentile using fraction 0.2, and global imputation. The data were then exported and further analyzed in RStudio using MSStats ^79^, removing sparse precursors and setting a cutoff for missing values at 0.5. The protein quantification and differential expression tables were used to generate volcano plots. Proteins were considered significantly enriched if they displayed a p-value < 0.1 (90% confidence over four biological replicates) and a log_2_FC ratio > 0.75 (168 % of protein enrichment). Before mass spectrometric analysis, the input, elute, and flow-through samples were run on Bio-Rad precast gradient gels and transferred at 100V for 2 hours. Membranes were blotted with streptavidin-HRP (1:2000) for 1 hour at room temperature and with anti-FLAG-M1 (1:1000) overnight at 4 °C.

### Statistical analysis

The statistical analyses were performed using GraphPad Prism (version 9). Experiments were carried out in triplicate, and all values were expressed as mean ± sem. Details of the statistical method applied and the number of data points (n = value, here “n” represents the number of organoids, slices, cells, or components) for each experiment are provided in the figure and its legend. Quantification graphs carry “n” and “p” values. In all cases, a p-value < 0.05 was considered statistically significant. Briefly, for two-group comparisons: Mann–Whitney U test, for three-group comparisons: Kruskal–Wallis test, followed by Dunn’s post-hoc test with Bonferroni correction, and for categorical proportions: Fisher’s exact test was performed. Differences were considered significant at p < 0.05. All tests were two-tailed.

### Data Availability

All reagents generated in this study are available from the Corresponding Author upon completion of a Materials Transfer Agreement. The raw single-cell RNA-seq data supporting the findings of this study have been deposited in the SRA under submission accession number PRJNAXXXX. The gene expression count matrix, extended metadata, and accompanying code notebook are available at Zenodo: https://doi.org/10.5281/zenodo.16738920. For further information and requests for resources and reagents, contact the lead, Jay Gopalakrishnan.

### Code Availability

The original code for the single-cell RNA-seq analysis of CCB79 brain organoids is publicly available on GitHub at https://github.com/CCBLAB-UKJ/CCB79_analysis_reproducibility and archived on Zenodo at https://doi.org/10.5281/zenodo.16738920.

---

**Supplementary Figure 1 (Related to main Figure 1).**
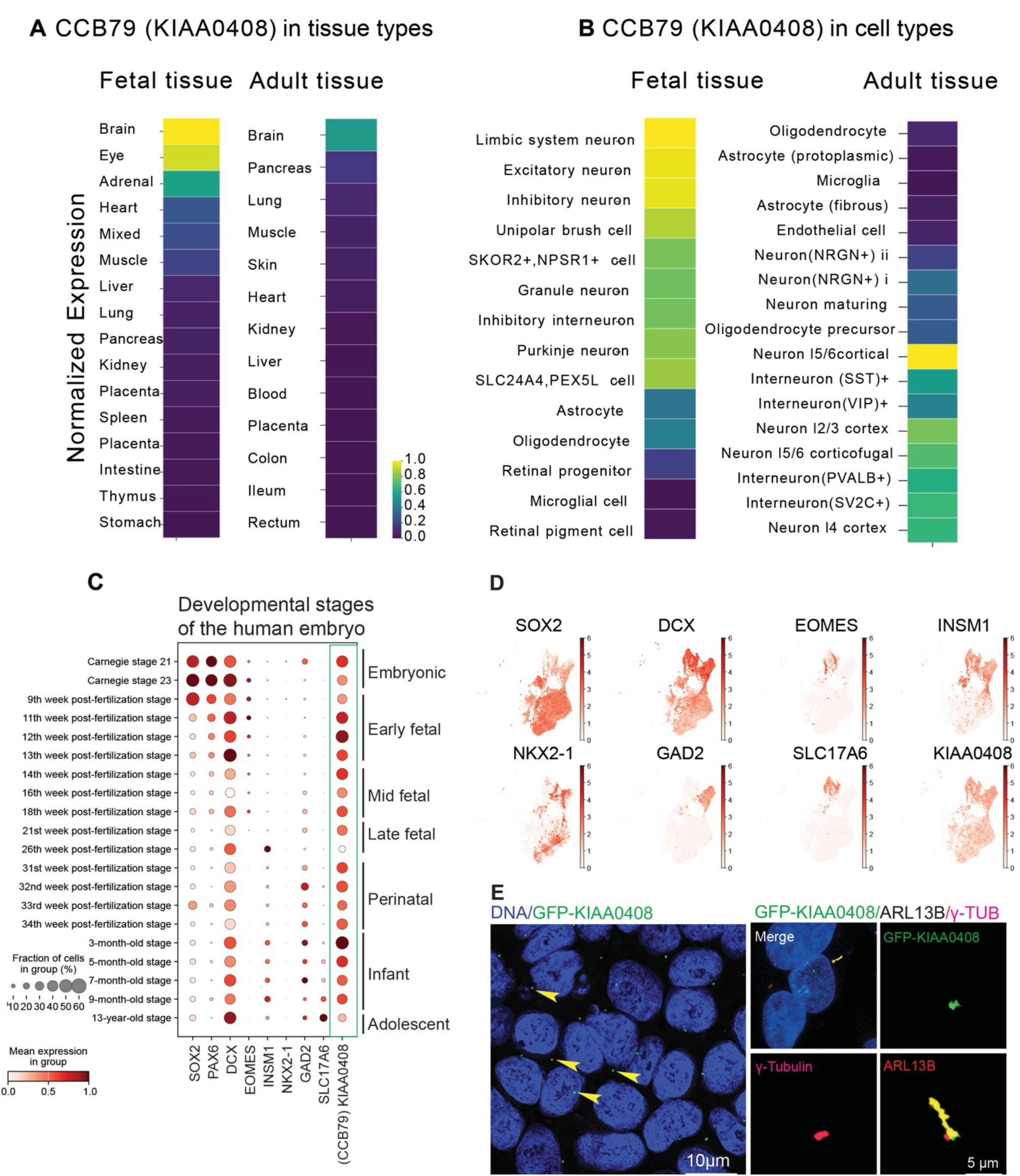
**A.** Heatmaps display the minimum–maximum normalized expression of CCB79 (*KIAA0408*) across multiple types of fetal (left) and adult (right) tissues, using GTEx V8 data. **B.** Expression of CCB79 (*KIAA0408*) across fetal and adult brain cell types, with all values normalized to the fetal expression range to emphasize developmental differences. **C.** The dot plot shows the standardized expression of key developmental and neuronal marker genes across 20 defined stages of human neocortical development. Dot size reflects the proportion of expressing cells in each group, while color indicates the normalized expression level. CCB79 (*KIAA0408*) expression is highlighted with a green box, indicating a similar expression profile to that of early neurons expressing DCX. **D.** UMAP showing distribution of SOX2, DCX, EOMES, INSM1, NKX2-2, GAD2, SLC17A6 and KIAA0408. **E.** GFP-KIAA0408 (green) expression in iPSCs showing a single dot indicated by yellow arrowheads. Scale bar 10 µm. Left panel: GFP-KIAA0408 (green) is at the base of the cilium (ARL13B, yellow). γ-tubulin labels the mother and daughter centrioles (red). Scale bar 5 µm.

**Supplementary Figure 2 (Related to main Figure 1 and 2).**
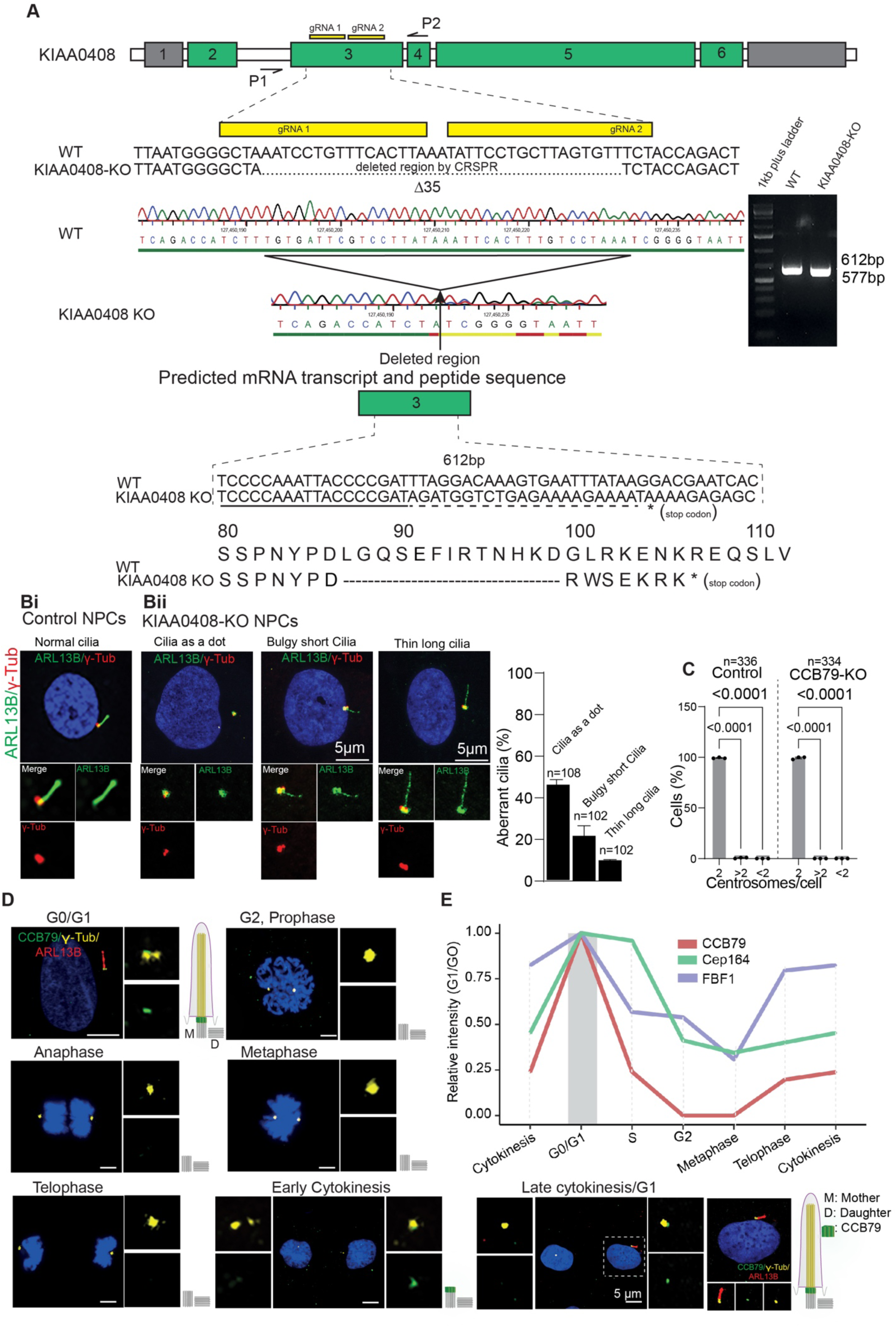
**A.** Exon structure of *KIAA0408*, positions of guide RNA to ablate the gene, and the results of gene ablation. The region deleted after CRISPR editing is shown. Agarose gel shows the fragment after the deletion. The predicted peptide with an induced stop codon is shown in the bottom panel. **B. (i)** Control NPCs show normally ciliated cells (ARL13B, green). γ-tubulin labels the mother and daughter centrioles (red). **(ii)** CCB79-KO NPCs show no ciliated cells. If cilia are observed, the following categories apply. No cilia, cilia as a dot, bulgy, short, or thin, long. (ARL13B, green). γ-tubulin labels the mother and daughter centrioles (red): scale bars 20 µm and 5 µm. The bar diagram at the bottom quantifies the various categories of cilia in CCB79-KO cells. Data are presented as mean ± SEM from at least three independent experiments. The “n” numbers are given in the graph. **C.** Centrosome numbers are unchanged in CCB79-KO NPCs. A bar diagram quantifies centrosome numbers between control and CCB79-KO cells. Data are presented as mean ± SEM from at least three independent experiments. A two-way ordinary ANOVA, followed by Sidak’s multiple comparisons test, was used. The “n” numbers and the “p” values are given in the graph. **D.** CCB79 staining across the cell cycle stages in NPCs. CCB79 (green) appears only at the start of ciliogenesis (ARL13B, red) at G0, disappears, and reappears in one of the late cytokinesis cells that is ciliated: γ-tubulin labels mother and daughter centrioles (yellow). The diagram of the mother (M) and daughter (D) centrioles and cilia is provided on the side to aid in interpreting the data. Scale bar 5 µm. **E.** Relative intensities of DAP across the cell cycle. Intensities were calculated relative to DAP intensities at the ciliary base in G1/G0 cells. Note that CCB79 peaks only at G0/G1 and then declines.

**Supplementary Figure 3 (Related to main Figure 2).**
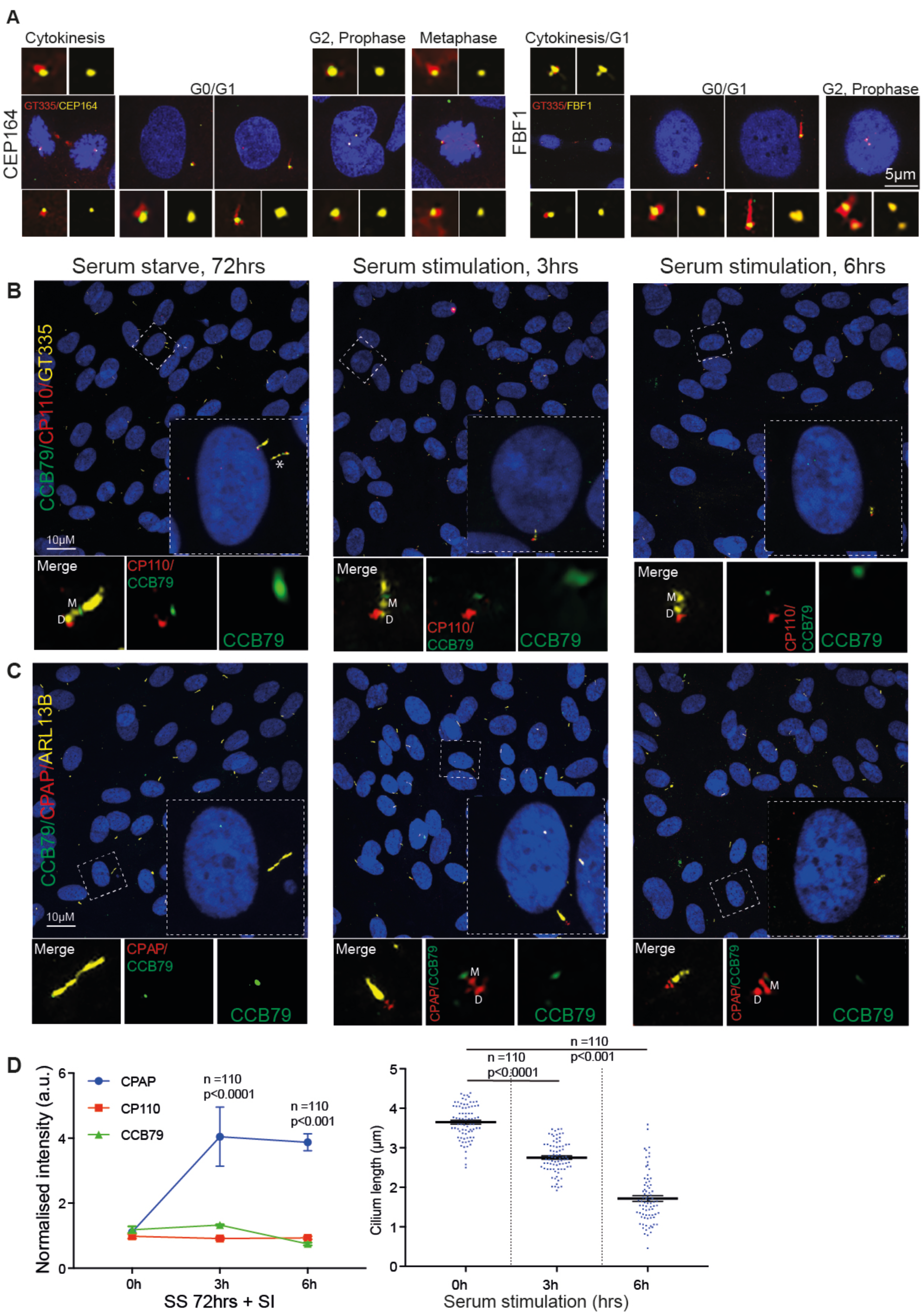
**A.** Localization behavior of DAP components, CEP164, and FBF1 (yellow) across various cell cycle stages. Cilia are labelled by GT335 (red). Note, in contrast to CCB79, which localizes only at the onset of cilium initiation (G0), these DAPs localize throughout the cell cycle stages. Scale bar 5 µm. **B.** Serum starvation and serum stimulation experiments using hTERT inactivated RPE1 cells. After 72 hrs of serum starvation, all cells are ciliated. CCB79 (green) is at the distal site of the mother centriole (M). Centrioles and cilia are labeled by GT335 (yellow). CP110 (red) labels only daughter centrioles (D). Cells begin to disassemble cilia after three hours of serum addition. Note that the CCB79 intensity is reduced (middle panel). The same observation is at six hours after serum addition. Dotted boxes indicate the cells that are zoomed in on the panel. Cilia are shown in the magnified panel. Asterik is not the cilium from the same cell. Scale bar 10 µm. **C.** The same experiment, which includes CPAP (red), a cilium disassembly factor, which, in contrast to CCB79, gains intensity. Scale bar 10 µm. **D.** Graphs that quantify the findings, such as intensity of CPAP (blue), CP110 (red), and CCB79 (green). A scatter plot displays the cilia length at various time points following serum addition. Data are presented as mean ± SEM from at least three independent experiments, and a non-parametric Mann-Whitney test was performed. The “n” numbers and p-values are provided in the graph.

**Supplementary Figure 4 (Related to main Figure 3).**
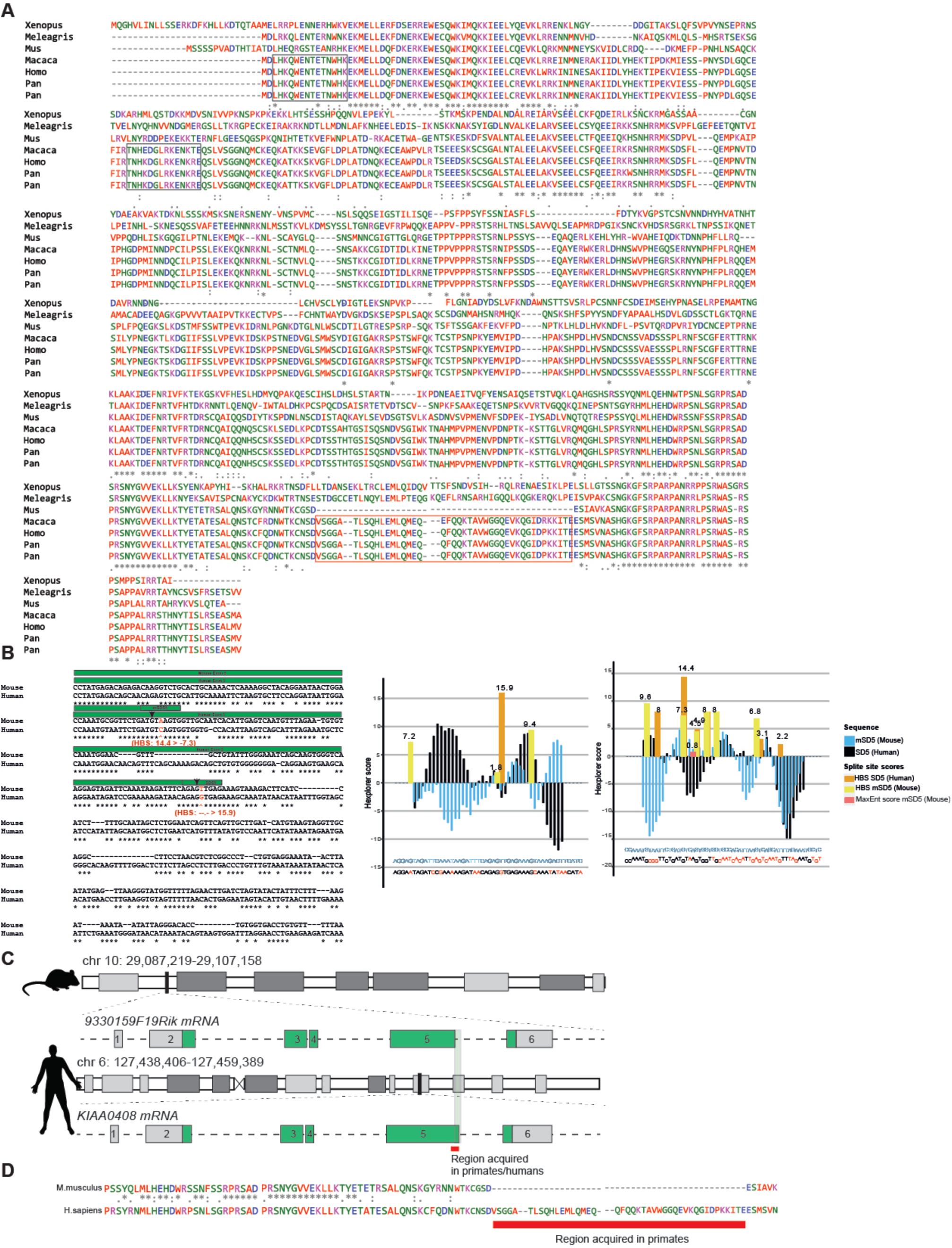
**A.** Pairwise alignment of amino acid sequences from different species is shown on the left. The boxed area is the epitope used to generate anti-CCB79 antibodies. The red boxed area, positioned at the end of exon five, indicates the region gained in primates, which comprises 43 amino acids. (Describe different species as abbreviations) **B.** Sequence alignment of the region, including exon 5, indicates changes in alternatively spliced exons between the mouse and human genomes. Bases in red with an arrow indicate the area where a primate-specific point mutation likely occurred in the evolutionary time between man and mouse. Asterisks indicate conserved positions between both species. Splice site strength and splicing regulatory landscape predictions using scoring algorithms (MaxEnt, H-bond score) were analyzed by Hexplorer. The HEXplorer score profiles represent splicing regulatory potential across the exon-intron boundary. The splicing region modified by the point mutation in humans exhibits a pronounced enhancer regulatory region and a more favorable splice site (splicing domain five or SD5), as indicated by a high HBS (15.9). This occurs downstream of the mouse splice site (mouse splicing domain five or mSD5), which is further suppressed by a second point mutation in humans (HBS score changes from 14.4 to −7.3), thus enabling exonization in humans and other primates. **C.** Gene structure and the coding regions of the mouse gene compared to human CCB79. The area gained in primates is highlighted. **D.** Mouse and human protein sequences are given. The red bar underlines the region of 43 amino acids that is unique to primates but is absent in the mouse protein.

**Supplementary Figure 5.**
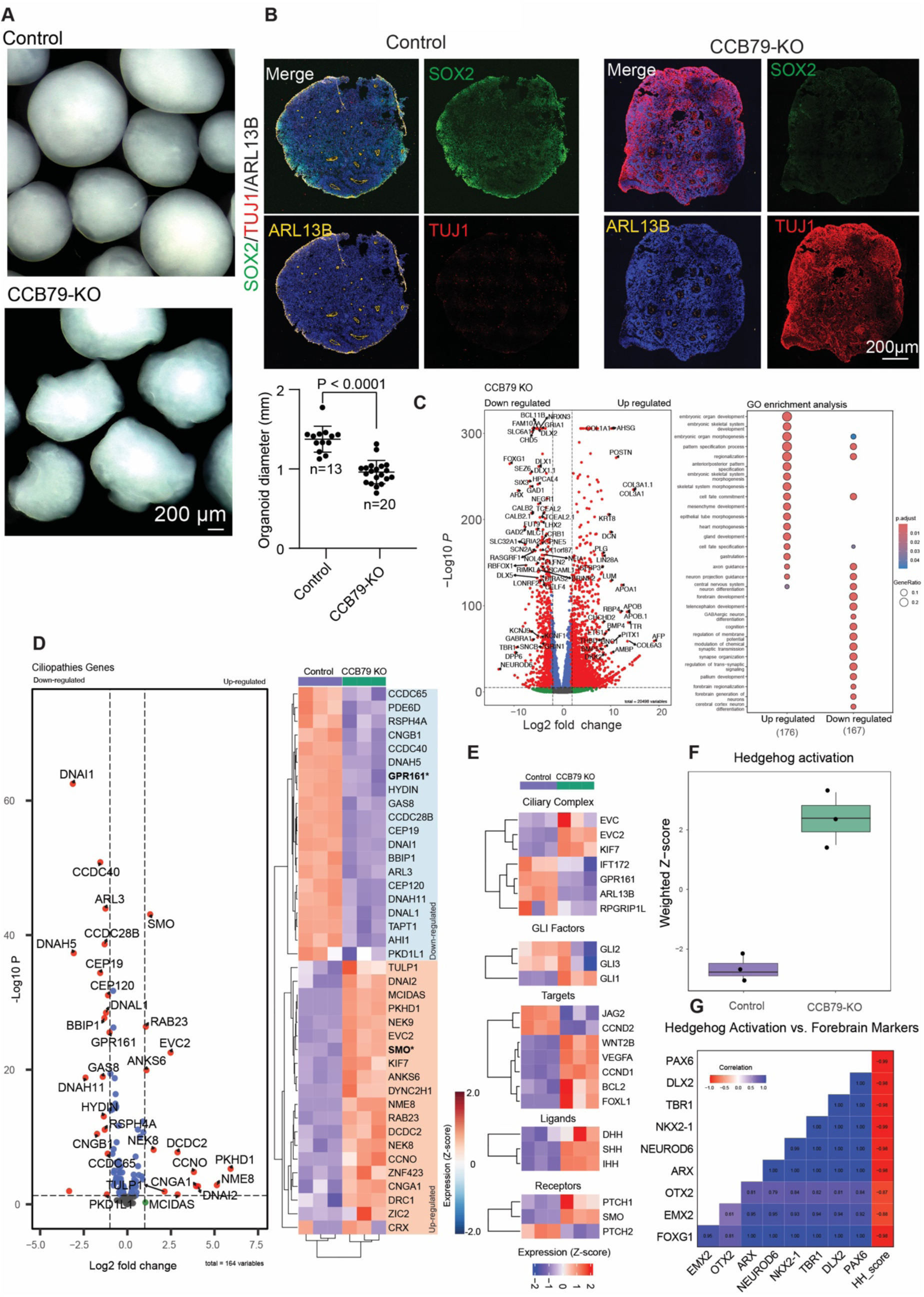
**A.** Macroscopic images of control and CCB79-KO brain organoids. Scale bar 200 µm. The scatter plot below quantifies individual brain organoids and shows that the CCB79-KO brain organoids are significantly smaller in size. **B.** Thin-sectioning and imaging of whole surface brain organoids. Control shows typical architecture with VZ, whose lumen is lined by cilia (ARL13B, yellow). SOX2 (green) shows NPCs. At this magnification, there are only barely visible TUJ1-positive neurons at the basal side (left panel). CCB79-KO organoids in the right panels show a disorganized cytoskeleton characterized by the loss of VZ, SOX2-positive progenitors with a gain of TUJ1-positive neurons all over the place. Scale bar 200 µm. **C.** Left panel: Volcano plot depicting differential gene expression with log2 fold-change on the x-axis and –log10 p-value on the y-axis. Significantly upregulated and downregulated genes (log_2_FC > 2, FDR < 0.05) are highlighted between the CCB79-KO and control samples. Right panel: Dot plots of enriched Gene Ontology Biological Processes (GO: BP) for the top 200 upregulated and downregulated genes in CCB79-KO. **D.** Left panel: Volcano plot illustrating differential gene expression, highlighting confirmed ciliopathy genes—right panel: Heatmap of the top 50 most variable ciliopathy genes, scaled by z-score. Columns represent samples grouped by control (purple) and CCB79-KO (green). **E.** Categorical heatmap of HH-related gene groups (Ligands, Receptors, GLI factors, Ciliary components, Suppressors, Targets). Columns represent samples grouped by control (purple) and CCB79-KO (green). **F.** Weighted HH signaling activity score computed by integrating activator, suppressor, and receptor gene expression z-scores shown as a boxplot comparing control and CCB79-KO. **G.** Correlation heatmap showing Pearson correlation coefficients between HH signaling activity scores and forebrain developmental markers (FOXG1, EMX1, PAX6, TBR1, ARX, NKX2-1, and NEUROD6), color-coded from blue (positive) to red (negative) correlation.

**Supplementary Figure 6 (related to the main Figure 5).**
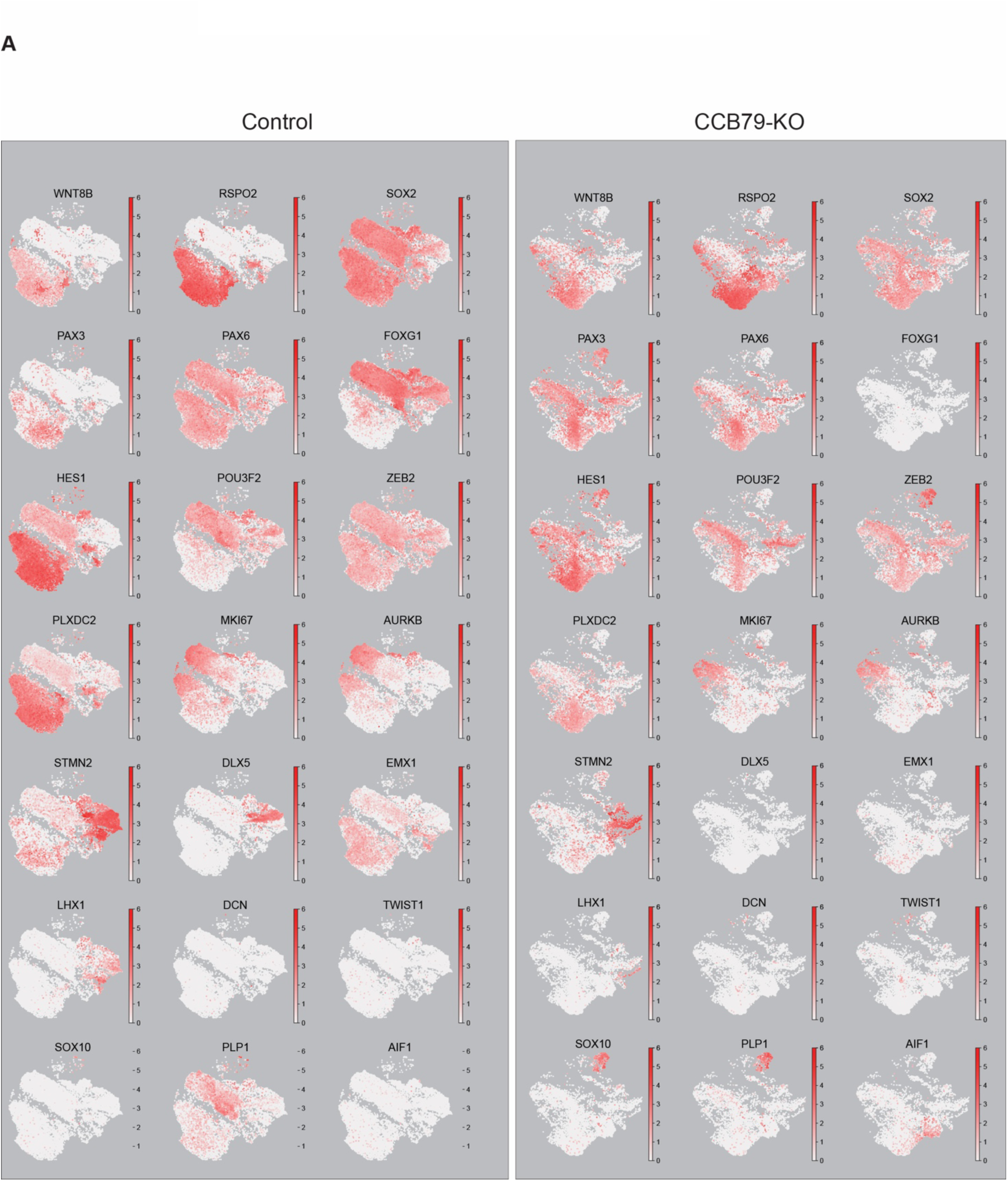
**A.** UMAP plots showing the distribution of various gene set markers used to annotate cell cluster types observed in control and CCB79-KO brain organoids.

**Supplementary Figure 7 (related to the main Figure 5).**
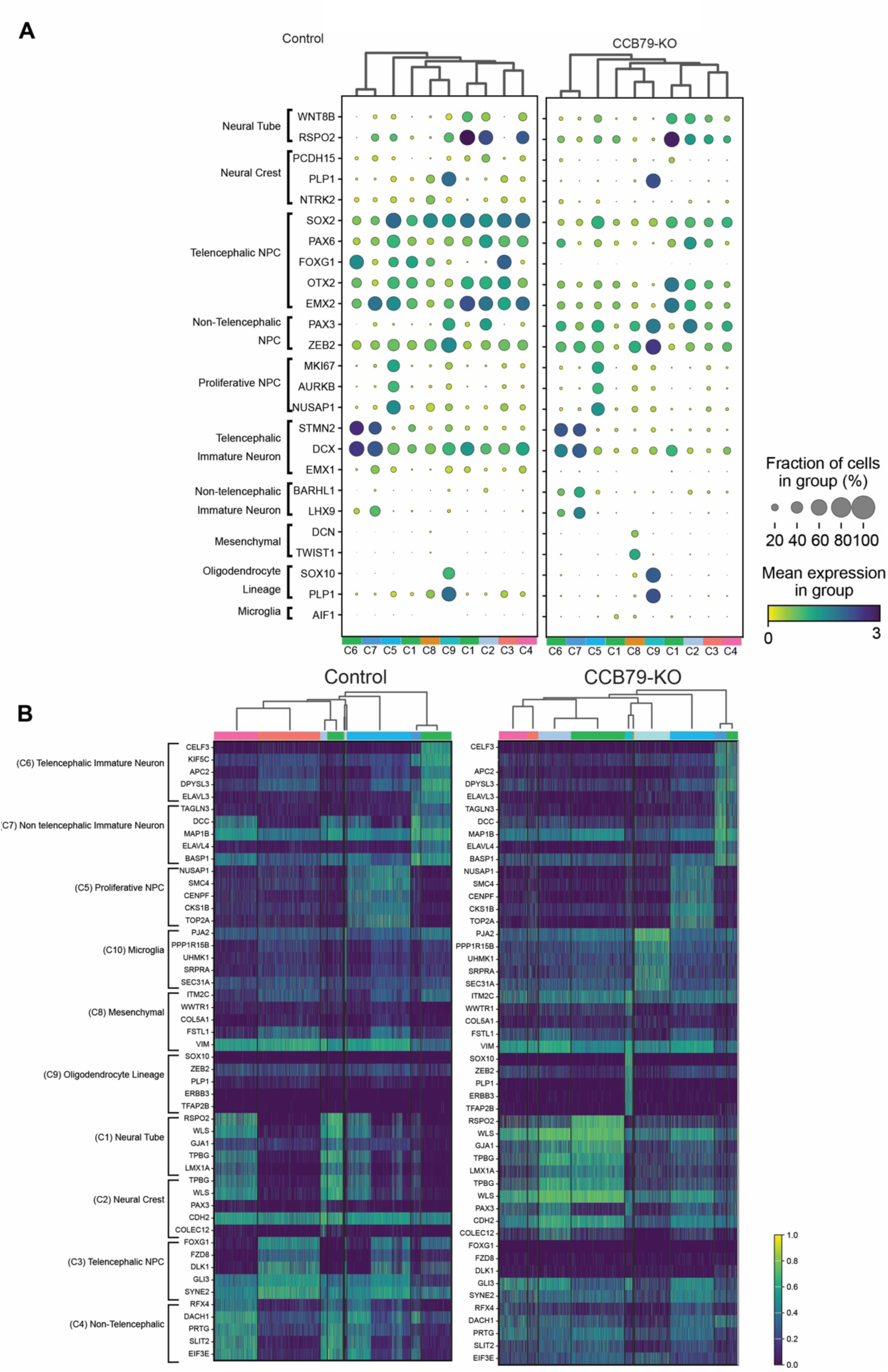
**A.** Dot plot showing expression of lineage- and region-specific markers across annotated cell types. Dot size represents the fraction of expressing cells; reversed viridis color intensity indicates the average scaled expression, ranging from 0 to 3. **B.** Hierarchical clustering represented as heatmaps showing the top 5 differentially expressed genes per cell type identified in control (left) and CCB79-KO (right) conditions. Color intensity reflects scaled expression, with gene names and corresponding cell types annotated.

**Supplementary Figure 8 (related to the main Figure 5).**
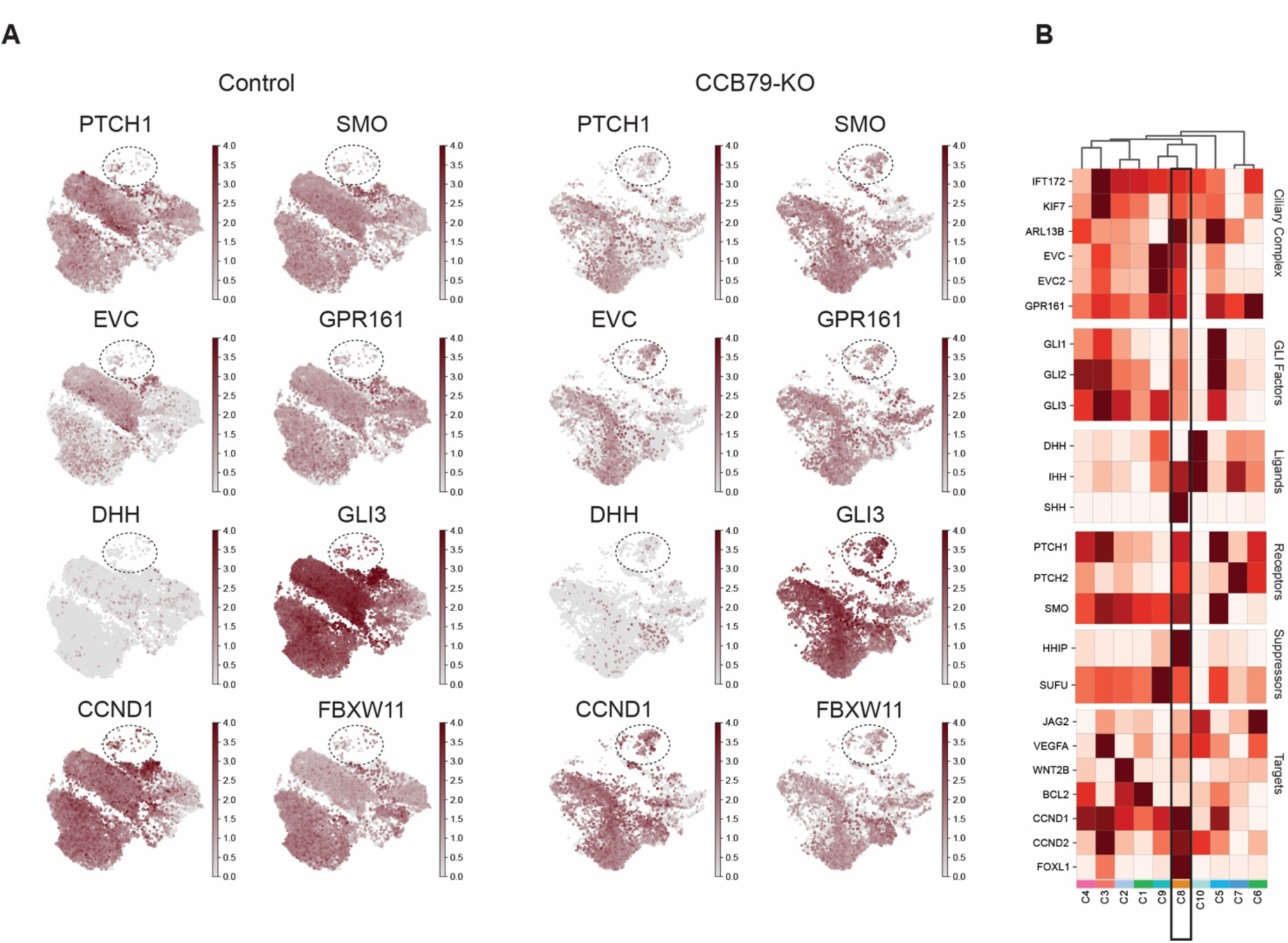
**A.** The UMAP showing the spatial distribution and normalized expression levels of the selected HH pathway in control (left) and CCB79-KO (right) conditions reveals striking differences in distribution, particularly in the oligodendrocyte precursor-containing cluster (dotted circle). Color intensity (gray-to-brown colormap) represents a scaled expression of 0–5. **B.** Matrix plots display the average expression (z-score) of HH pathway genes grouped by functional categories across annotated cell types. Rows represent HH pathway gene sets (Ciliary Complex, GLI Factors, Ligands, Receptors, Suppressors, Targets), and columns represent cell clusters. Dendrograms display the hierarchical clustering of genes and cell types based on their expression similarity. Color scale ranges from low (light pink) to high (dark red) expression. The most affected cluster 9 is highlighted with a box.

**Supplementary Figure 9 (related to the main Figure 5).**
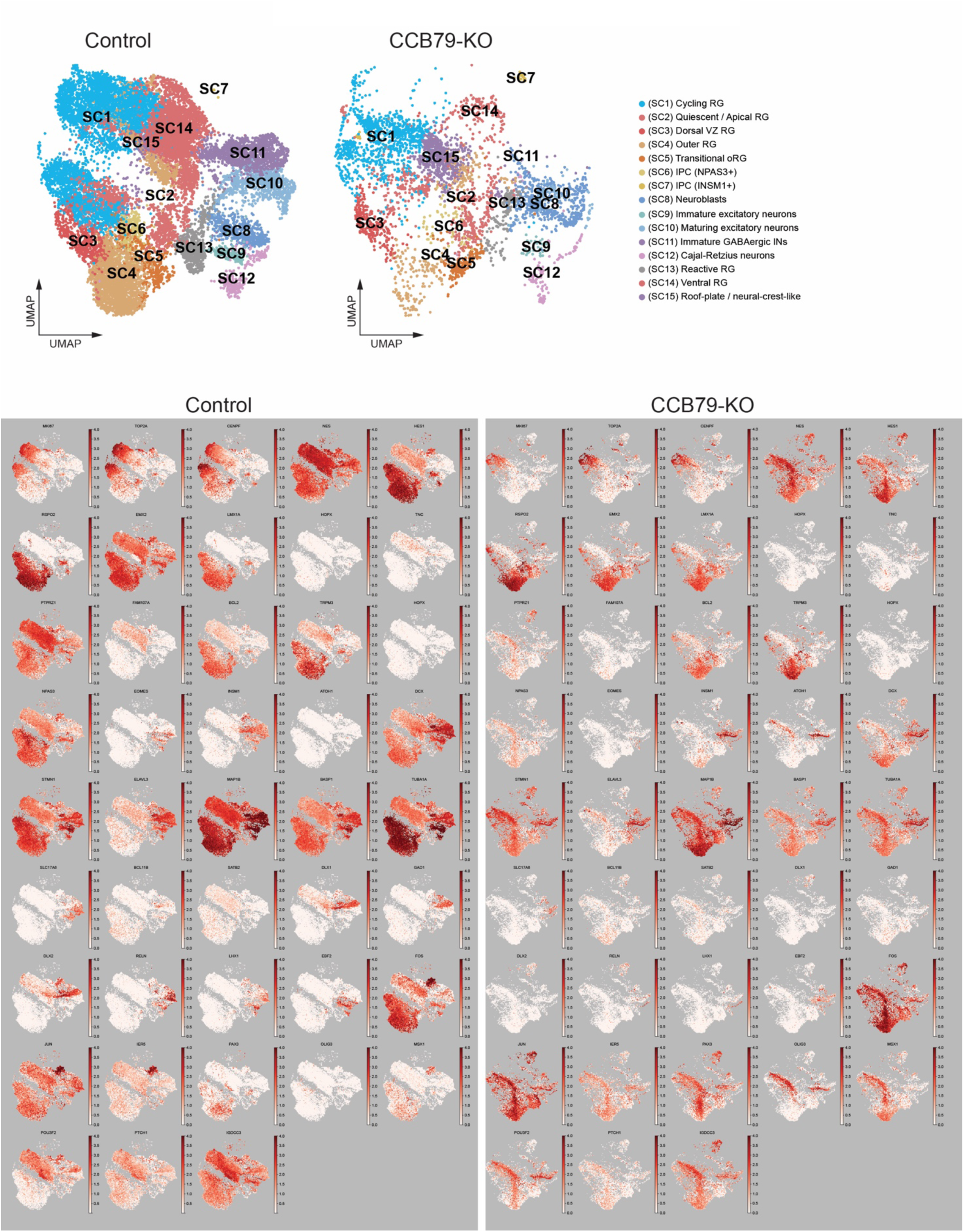
**A.** UMAP embedding of all cells colored by 15 major developmental cell type clusters after sub-clustering in control (left) and CCB79-KO (right) organoids. Legends represent colors that correspond to annotated cell types. **B.** UMAP plots showing cell type-specific expression of selected markers used to assign cell types in control (left) and CCB79-KO (right).

**Supplementary Figure 10 (related to the main Figure 5).**
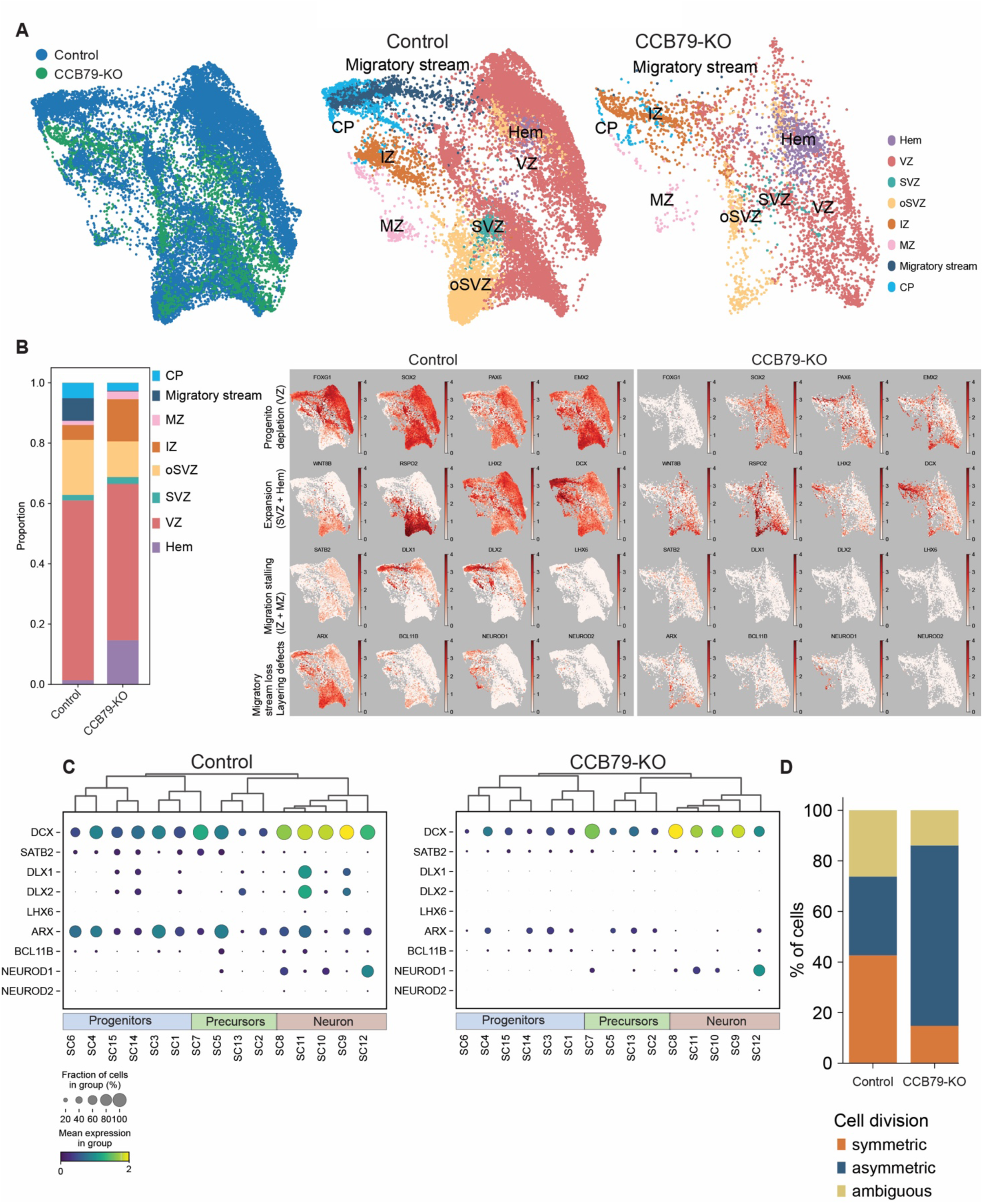
**A.** Right panel, Force-directed graph (FA layout) showing combined developmental trajectories, control = green, CCB79-KO = purple. Left panel, annotated regions Hem (lavender), VZ (red), SVZ (orange), oSVZ (mustard yellow), IZ (beige), MZ (light pink), Migratory stream (pink), CP (sky blue), comparing control vs CCB79-KO. **B.** Stacked bar plot of proportional representation of cells across ordered cortical layers comparing control vs CCB79-KO. **C.** Dot plot quantifies the relative expression of markers (left side) in progenitors, precursors, and neurons between control and CCB-KO brain organoids. **D.** Classification of progenitor division modes using gene module scoring for symmetric and asymmetric division markers. The bar plot displays the stacked percentages for symmetric, asymmetric, and ambiguous cases. Each panel illustrates distinct aspects of cortical development and perturbations in CCB79-KO organoids, including regional mispatterning, altered layer composition, and division-type imbalance.

**Supplementary Figure 11 (related to the main Figure 4,5 and 7).**
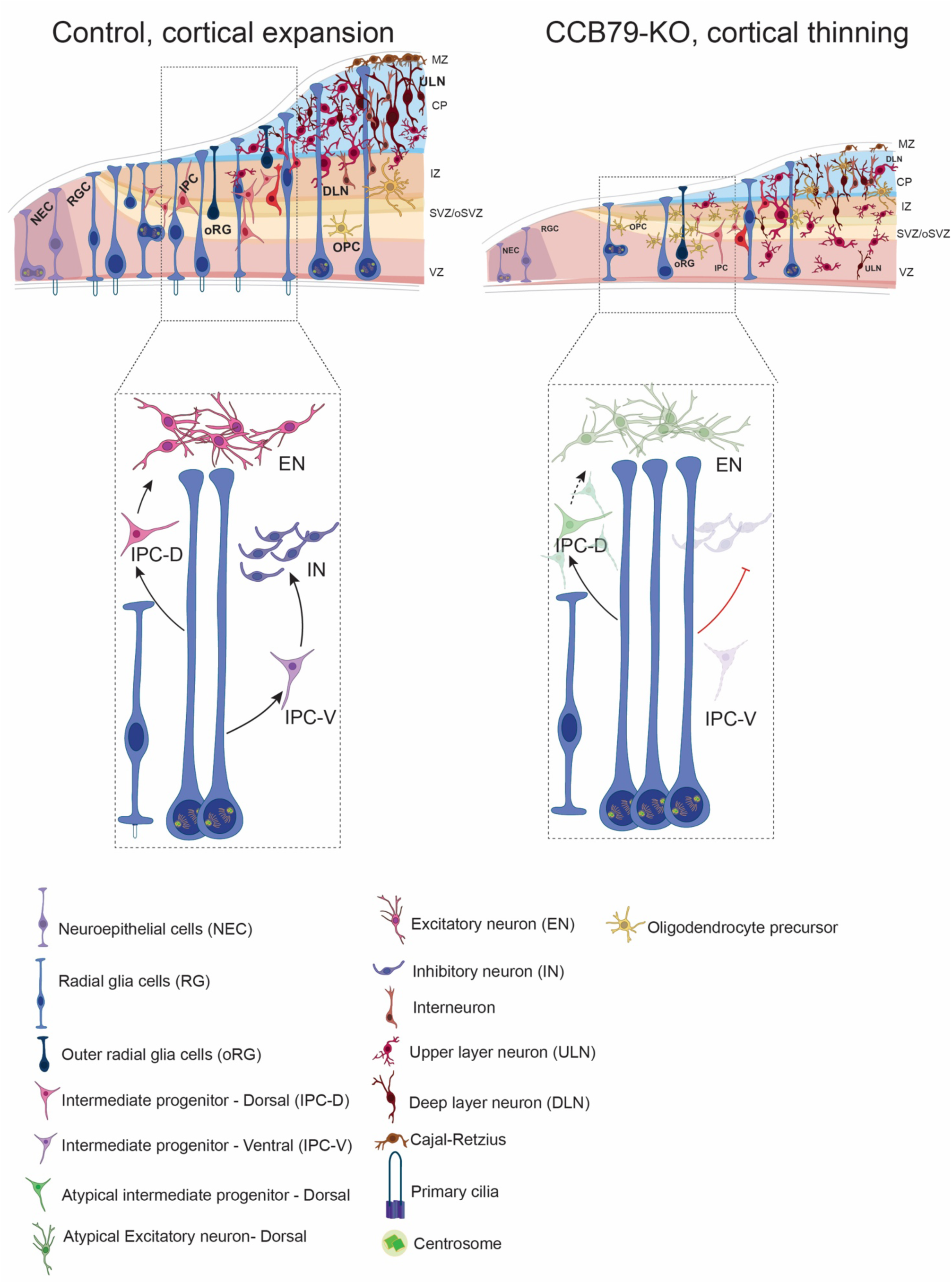
A schematic representation of the normal brain development (Left) compared to the absence of CCB79 (Right). In the absence of CCB79, primary cilia fail to assemble, which results in abnormal cell fate determination. CCB79-KO brain organoids have aberrant levels of oligodendrocyte precursor cells (OPC) at the expense of ventral progenitors, and the developing cortex is thinner than the control organoids with a decline in radial glial cells. Zoomed in panels: In the absence of CCB79, IPC-V (Ventral intermediate progenitors) are affected, and thus, no inhibitory neurons are produced. In the case of the dorsal pathway, the precursors do not display typical dorsal patterns (IPC-D) or populations. They are instead stalled in their migratory route, resulting in an aberrant level of excitatory neurons. The legends below show different cell types, centrosomes, and cilia.

**Movies 1 and 2.**

Volume rendered images of representative VZs from panel A for 3D volumetric reconstruction in control (Control movie 1) and CCB79-KO brain organoids (CCB79-KO movie 1). Rendering was performed using Arivis Pro. A close-up view offers a tunnel-traveling effect for visualizing the progenitor organization and its cilia at the apical region.

## Tables

**Supplementary Table 1**

Bulk RNA-seq differential expression analysis and Gene Ontology (GO) enrichment results.

**Supplementary Table 2**

Processed mass spectrometry data from miniTurbo-based proximity labeling experiments. Sheet 2 contains curated list of proteins with proximity network of CCB79.

**Supplementary Table 3**

Top 50 marker genes for each cluster identified in the integrated single-cell RNA-seq analysis.

